# Host Phylogeny Shapes Viral Transmission Networks in an Island Ecosystem

**DOI:** 10.1101/2022.10.04.510907

**Authors:** Rebecca K. French, Sandra H. Anderson, Kristal E. Cain, Terry C. Greene, Maria Minor, Colin M. Miskelly, Jose M. Montoya, Chris G. Muller, Michael W. Taylor, Kākāpō Recovery Team, Edward C. Holmes

## Abstract

Viral transmission between host species underpins disease emergence. Both host phylogenetic relatedness and aspects of their ecology, such as species interactions and predator-prey relationships, may govern cross-species virus transmission and zoonotic risk, although their relative impact is unknown. By characterising the virome of a relatively isolated island ecological community linked through a food web we show that phylogenetic barriers result in distantly related host species sharing fewer viruses. Host ecology had a much smaller influence on overall virome composition. Network analysis revealed that hosts with a high diversity of viruses were more likely to gain new viruses, and that generalist viruses were more likely to infect new hosts. Such a highly connected ecological community heightens the risk of disease emergence, particularly among closely related species.

**One-Sentence Summary:** Sequencing of an entire island virome reveals that closely related hosts have highly connected virus communities, increasing emergence risk.

---

Determining how viruses move through ecosystems is central to understanding their occasional emergence as pathogens. Cross-species virus transmission is a near universal feature of viruses (*1*). However, most studies of virus host jumping have focused on single virus pathogens and/or single host species (*2, 3*), limiting our capacity to reveal broad-scale host-virus interactions (*4, 5*). In addition, recent metagenomic studies suggest that only a tiny proportion of viruses cause disease, with apparently healthy wildlife species commonly infected with multiple viruses (*6*). In reality, viruses are a key component of global ecosystems, regularly moving between species in the absence of overt disease (*7*). A full understanding of infectious disease emergence therefore requires an ecosystem-level approach (*7, 8*), in which the viromes of entire interacting communities are investigated.

Two factors have been proposed to govern virus movement from one host species to another, shaping similarities and differences in virus composition among taxa, and hence determining virome structure at the ecosystem scale. First, it is possible that host phylogenetic relatedness directly impacts the frequency and pattern of cross-species virus transmission. Rates of cross-species virus transmission are expected to be higher among closely related host species, reflecting a greater similarity in the virus and host proteins that underpin successful virus-cell relationships such as host cell binding (*9*). A fundamental difference in virus-host cell relationships explains why viruses from vertebrates and invertebrates are usually phylogenetically distinct (*10*), even though the latter are often dietary components of the former. A second proposed factor is that ecological properties of the host play a major role in virus movement between host taxa by determining the probability of virus exposure (*11*). Each time species interact they provide opportunities for cross-species virus transmission. Hence, the more interactions, the greater the probability of host jumping. For example, changes in land use have increased human–animal interactions, driving disease emergence events in humans (*12*). Predator-prey interactions are common exposure events, with the consumption of prey providing direct exposure to viruses during digestion. Consequently, the structure of an ecosystem food web may have a large impact on the flow of viruses through communities. With the exception of marine microbial food webs in which viral lysis of microbial hosts impacts food web structure (*13*), this prediction is yet to be tested.

Metagenomic sequencing allows virus diversity to be explored with precision, by – in theory – sequencing the entire virome of samples without bias (*6*) and leading to the increasingly rapid discovery of novel viruses (*14*). Far less attention has been directed toward understanding how viruses are embedded within ecosystems, and the relative role of host phylogeny and ecology in structuring virus diversity. Research on food webs including viruses has commonly either focused on single pathogens of a species of interest (*15*), or on the microbial subset of the food web (*13*). The virome of an entire food web is yet to be characterised, in part because the large number of species in most ecological communities makes this impractical. However, in small island forest ecosystems, such as Pukenui/Anchor Island in the Fiordland region of southwestern New Zealand (fig. S1), the entire forest community is small enough that most broad taxonomic groups can be sampled, providing a snapshot of a food web virome.

The unique evolution of New Zealand wildlife (i.e., in the almost complete absence of native terrestrial mammals) means that unlike other forest ecosystems, the Pukenui/Anchor Island ecological community is dominated by birds, with no terrestrial mammals, and only one species of reptile. However, despite being small, isolated and having a low diversity (relative to forest ecosystems outside New Zealand), this community contains a breadth of ecological niches, including a variety of diets: carnivores, insectivores, plant-eaters (including herbivores, frugivores, nectarivores and granivores), piscivores and omnivores. This ecological community also contains numerous threatened species that may be vulnerable to disease emergence via cross-species transmission; for example, the critically endangered kākāpō (*Strigops habroptila*), endangered mohua (*Mohoua ochrocephala*), and critically endangered Te Kakahu/Chalky Island skink (*Oligosoma tekakahu*). The Pukenui/Anchor Island community also has a relative decoupling of phylogenetic relatedness and ecological niches. Some species that are distantly related (i.e., from different phylogenetic orders) possess a similar ecological niche and regularly interact. For example, yellow-crowned parakeets (*Cyanoramphus auriceps*, parrot) were frequently observed with passerines (brown creeper *Mohoua novaeseelandiae*, mohua, and the grey warbler *Gerygone igata*), travelling and foraging in multi-species flocks (pers. obs), which has been observed elsewhere in New Zealand (*16*).

We used metatranscriptomic (i.e., total RNA) sequencing to document the virome of each host in the community, including those that directly infect the host, bacteriophages, and viruses present in the host diet. Hence, we use the total viral diversity of each host, rather than only those viruses that have established a true infection, to provide a broader view of virus transmission. If host phylogeny were the key driver of viral diversity we would expect viromes to cluster according to the major host phyla (e.g., Chordata, Arthropoda and Streptophyta), as well as grouping at lower phylogenetic levels, with more closely related hosts having more similar viromes than hosts that are more distantly related (Fig. 1A). In contrast, if host ecology were the main driver of virus diversity, we would expect viromes to cluster according to the major dietary associations, with hosts that have similar diets possessing similar viromes, and predators and prey clustering together (Fig. 1B).

**Fig. 1.**
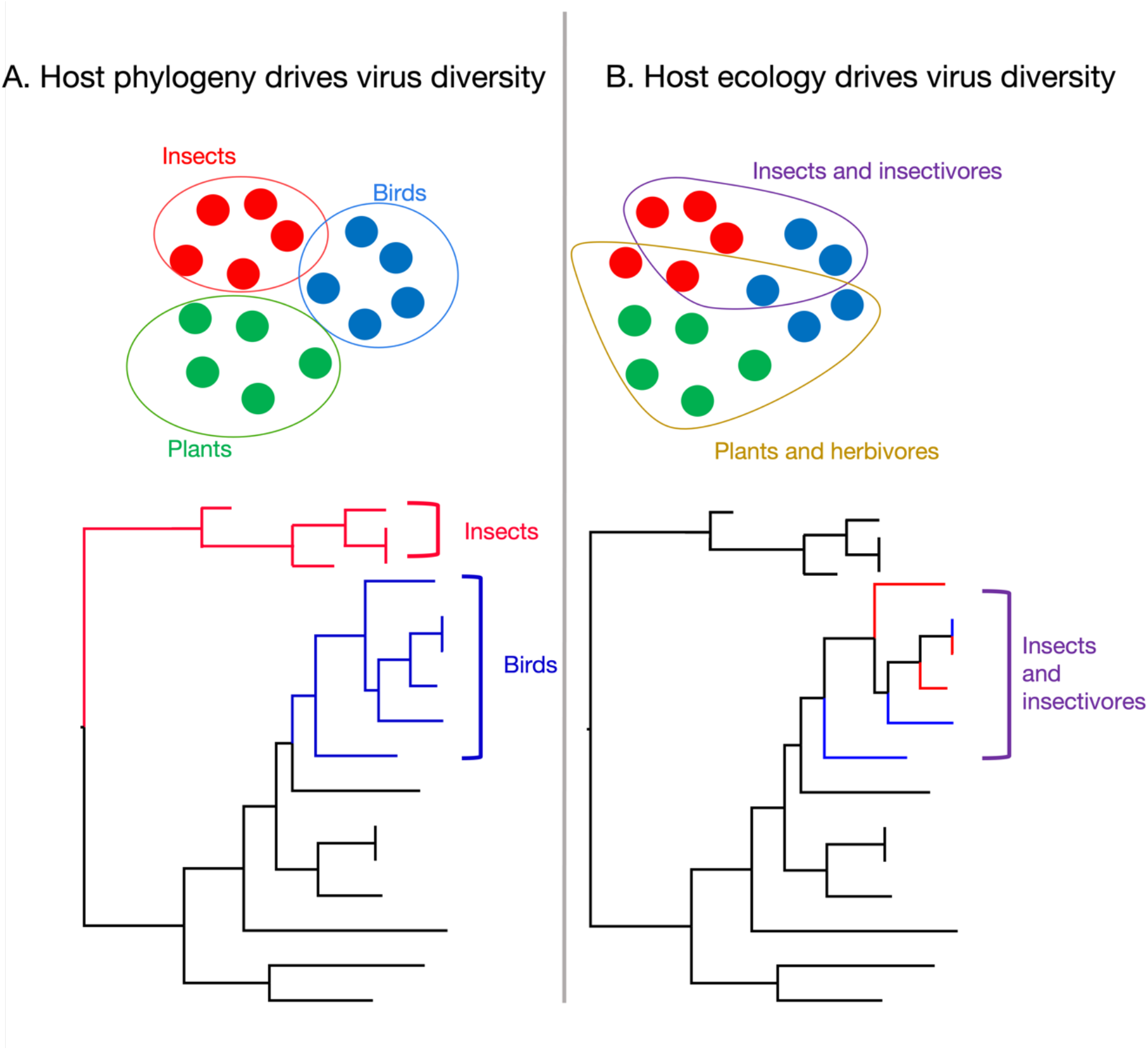
Expected impact of the two drivers of viral diversity on the structure of virus phylogenies. (**A**) Host evolutionary relationships and (**B**) host feeding ecology. Circles denote hypothetical clusters of hosts with similar viromes: i.e., dots (hosts) within a circle have more viruses in common than to other dots outside the circle.

Our sampling of the Pukenui/Anchor Island forest community included all key vertebrate species in addition to representative sampling of invertebrates and plants (table S1). The sampled community comprised five host phyla, 13 classes and 37 orders. We conducted diversity and network analyses to determine the importance of host phylogeny (at the phylum, class and order level) and host ecology on viral diversity at the virus family level. A detailed account of the methods used in this study is provided in the supplementary materials.

## Results

### Viruses predominantly cluster according to host phylogeny

We identified 16,633 viral sequences (assembled contigs) with an abundance of 360 million reads from a total of 112 different viral families. Non-metric multidimensional scaling plots revealed that, overall, viral communities clustered to the level of host phyla (Fig. 2), which was confirmed using pairwise adonis tests (table S2). All comparisons between Chordata, Arthropoda and Streptophyta were significant (p<0.05, n=49, table S2). In addition, host phylogenetic order explained the most variation between viral communities (R^2^=0.8), compared to host phylum (R^2^=0.13) and class (R^2^=0.30), when used as the dependant variable in the adonis test (n=49).

**Fig. 2.**
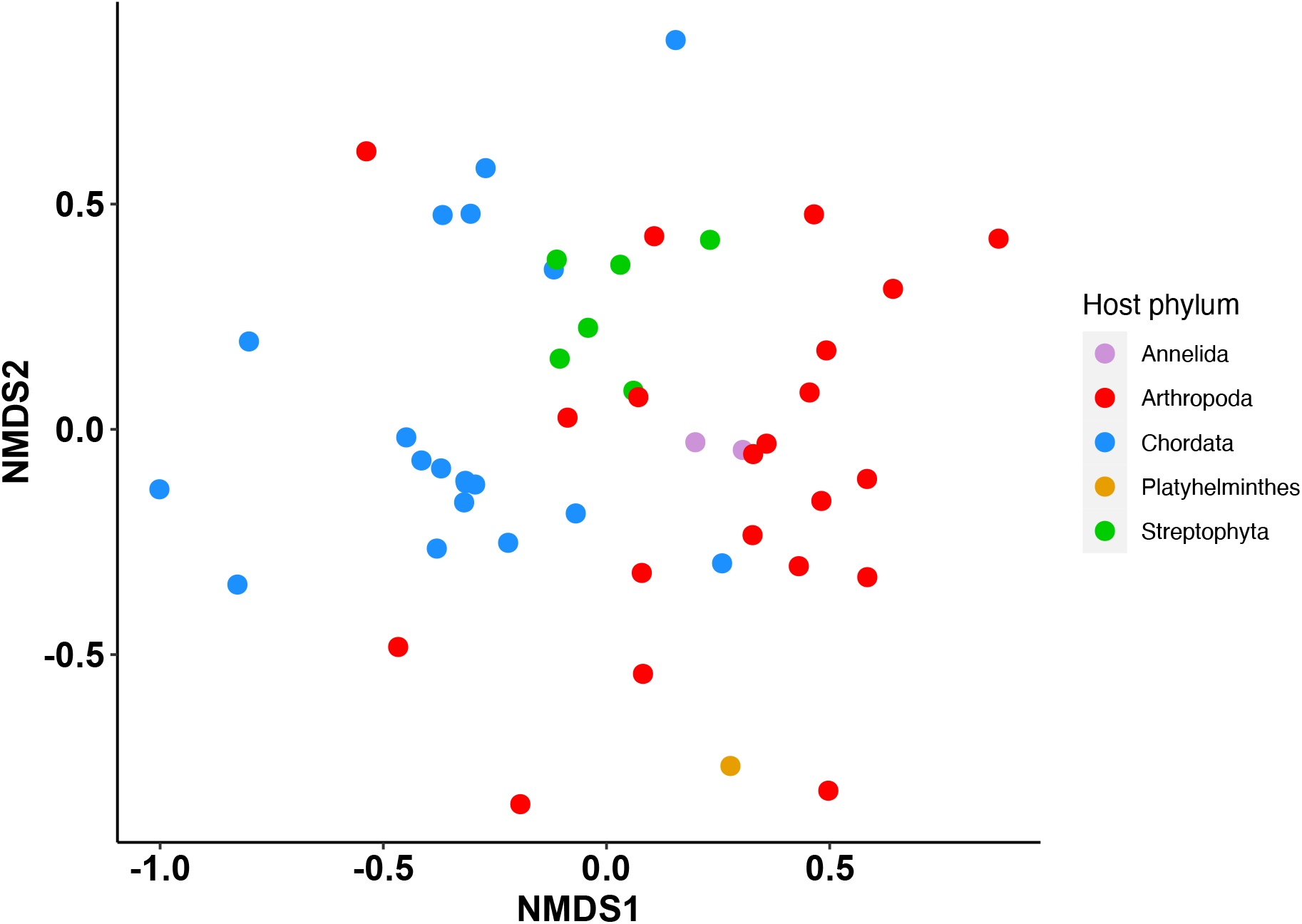
The similarity/dissimilarity of viral communities among host species. The first two dimensions of a three-dimensional non-metric multidimensional scaling (NMDS) plot show that virus communities at the family level cluster according to host phyla. Axes refer to the dimensions of the NMDS, stress = 0.196, n=49.

Within Chordata, we used host order and host diet to compare how much variation in the viromes is explained by each factor. Host diet was described using two separate binary factors based on the primary diet types (table S1) – ‘insectivore’ (yes/no) and ‘plant-eater’ (yes/no). Nectar, fruit and seed eaters were assigned as plant-eaters. Omnivores that feed on both invertebrates and plants were assigned as both an insectivore and a plant-eater. Carnivores and piscivores were assigned as neither an insectivore nor a plant-eater. When controlling for host order, permutational multivariate analysis of variance (PERMANOVA) tests revealed that insectivores had significantly different viromes than non-insectivores, and plant-eaters had significantly different viromes from non-plant-eaters (p<0.05, n=19, table S2). However, host order explained much more of the variation (R^2^=0.47) than diet (insectivore R^2^=0.06, plant-eater R^2^=0.07). A similar result was obtained when including only viruses likely to infect chordates, with all comparisons significant (p<0.05, n=19, table S2), and host order explaining more variation (R^2^ = 0.48) than host diet (insectivore R^2^=0.06, plant-eater R^2^=0.06).

To further explore this clustering of viral communities by host phylogeny, we created a bipartite network (herein referred to as the ‘host-virome network’) and identified four modules using a community detection algorithm. Notably, the modules follow host phylogenetic groupings, with a Pearson’s chi-squared test of independence indicating that host phylum and module were highly correlated (χ^2^ = 68.4, df = 12, p-value = 6.4 × 10^8^). Module one contained only invertebrates (arthropods, annelids and a platyhelminth), module two predominately comprised plants (75% plants and 25% invertebrates), while modules three and four largely comprised chordates (86 and 81% chordates, 14 and 19% invertebrates, respectively) (Fig. 3). All modules contained both DNA and RNA viruses. Module three comprised mostly bacteriophage, while the other modules were dominated by viruses that infect the hosts in those modules.

**Fig. 3.**
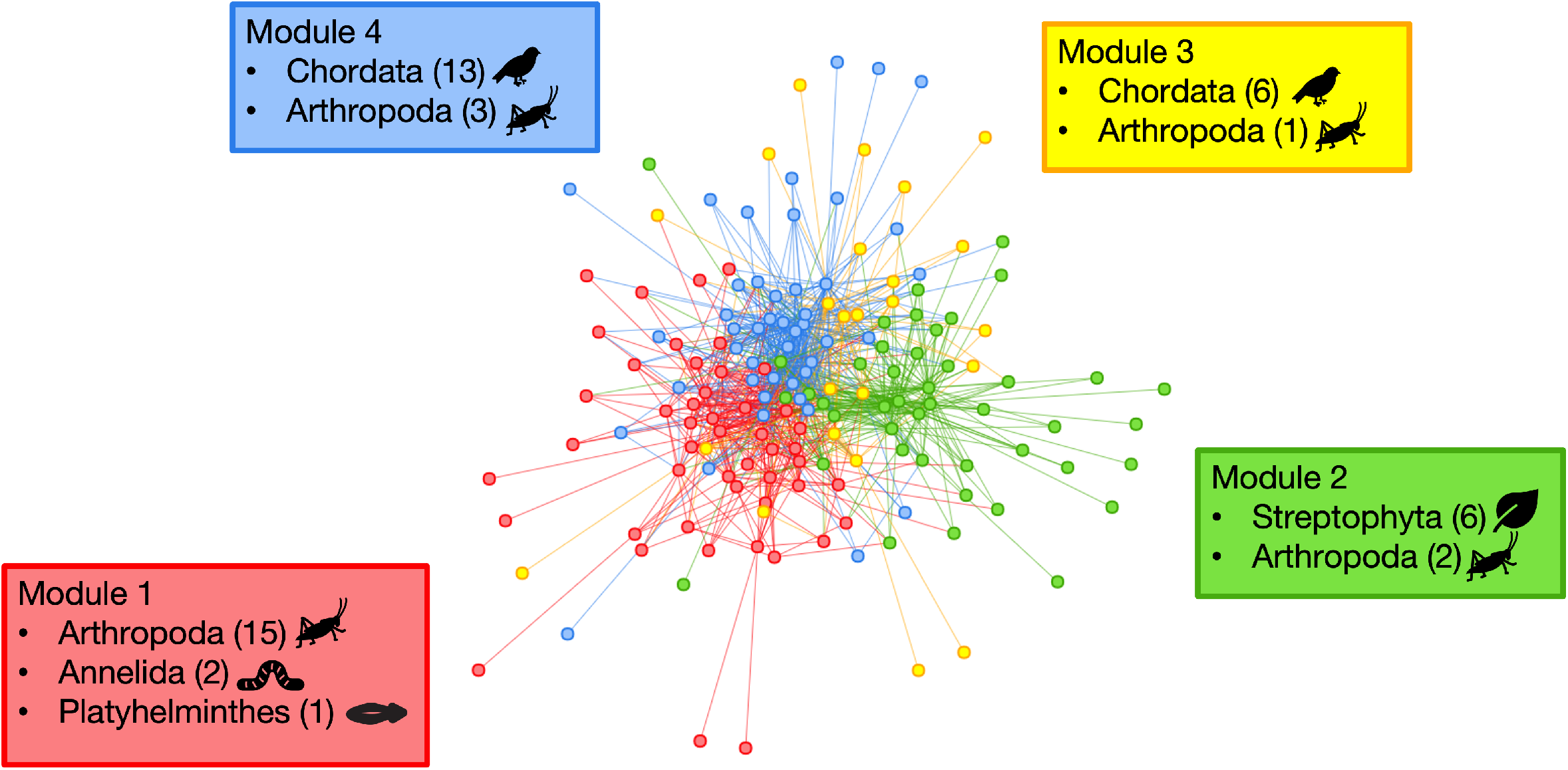
Host-virome network (bipartite) displayed using the Fruchterman-Reingold layout. The modules are shown by node colour. Nodes include both host library and virus families. The boxes show the host makeup of each module, with the number of hosts belonging to each host phylum shown in parenthesis.

Modules also had differing levels of host and virus richness, with module one containing the highest number of hosts (18), module two containing the highest number of virus families (36), and module three containing the lowest number of both hosts (7) and virus families (17). The communities within each module were significantly different from one another (p<0.05, n=49, pairwise adonis test; table S2), which was robust to rarefication (table S3). Interestingly, the two modules containing chordates (modules three and four) had the smallest difference in communities (table S2), although still statistically significant.

### Network structure

We next analysed the structure of the host-virome network using the degree distribution (i.e., the distribution of the number of links between nodes) in comparison to a network generated with a null model (a bipartite network with the same number of nodes and links, randomly assigned). The cumulative degree distributions for the host and virus nodes followed a truncated power-law distribution (Kolmogorov-Smirnov test p>0.05) of 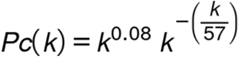 for the hosts and 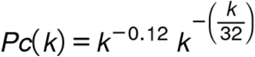 for the virus families, as shown by the fit lines in Fig. 4. The null networks also follow truncated power-law distributions of 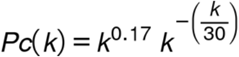 and 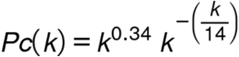, but with lower cut-off values than the host-virome network. Viruses with few connections had fewer than the random expectation, while viruses with more connections had more than expected by chance, shown by the null and virus distributions intersecting at approximately 10 links (Fig. 4). In contrast, all hosts had systematically more connections than the random expectation.

**Fig. 4.**
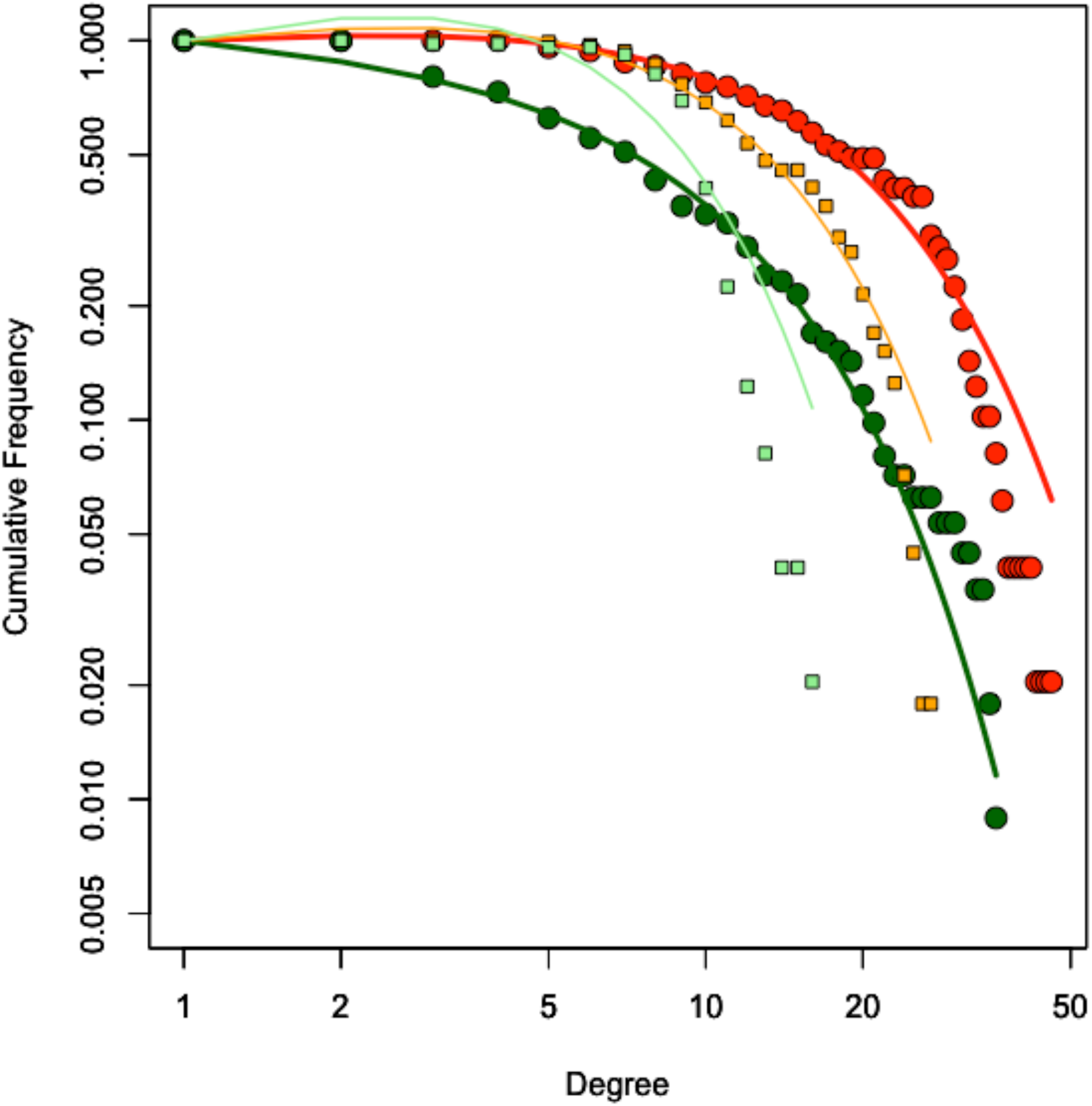
The degree distribution of the host-virome network (large circles) and the null network (small squares), shown as the cumulative probability of finding a virus family in the network with k or less-associated hosts (Pc(k)). The colour denotes node type, with red and orange referring to host nodes and the dark and light green the virus family nodes, in the host-virome network and null network, respectively.

### Potential for virus spread within the network

To examine the strength of connections between hosts that shared viruses, we created a unipartite network (i.e., links between hosts with shared virus families, where a link is a connection between two hosts) based on the Bray-Curtis dissimilarity matrix, which we refer to here as the ‘host community network’ (Fig. 5). This network had a high level of connectivity between hosts and within host phyla, with the connections across host phyla generally weaker (i.e., a higher Bray-Curtis value). The maximum shortest path was eight, such that the most distantly connected nodes were still only eight links (connections between hosts) away from one another. The mean shortest path distance was only 3.19, such that on average each node is 3-4 links away from every other node. Also of note is a key cluster containing predominately chordates with a high number of strong connections.

**Fig. 5.**
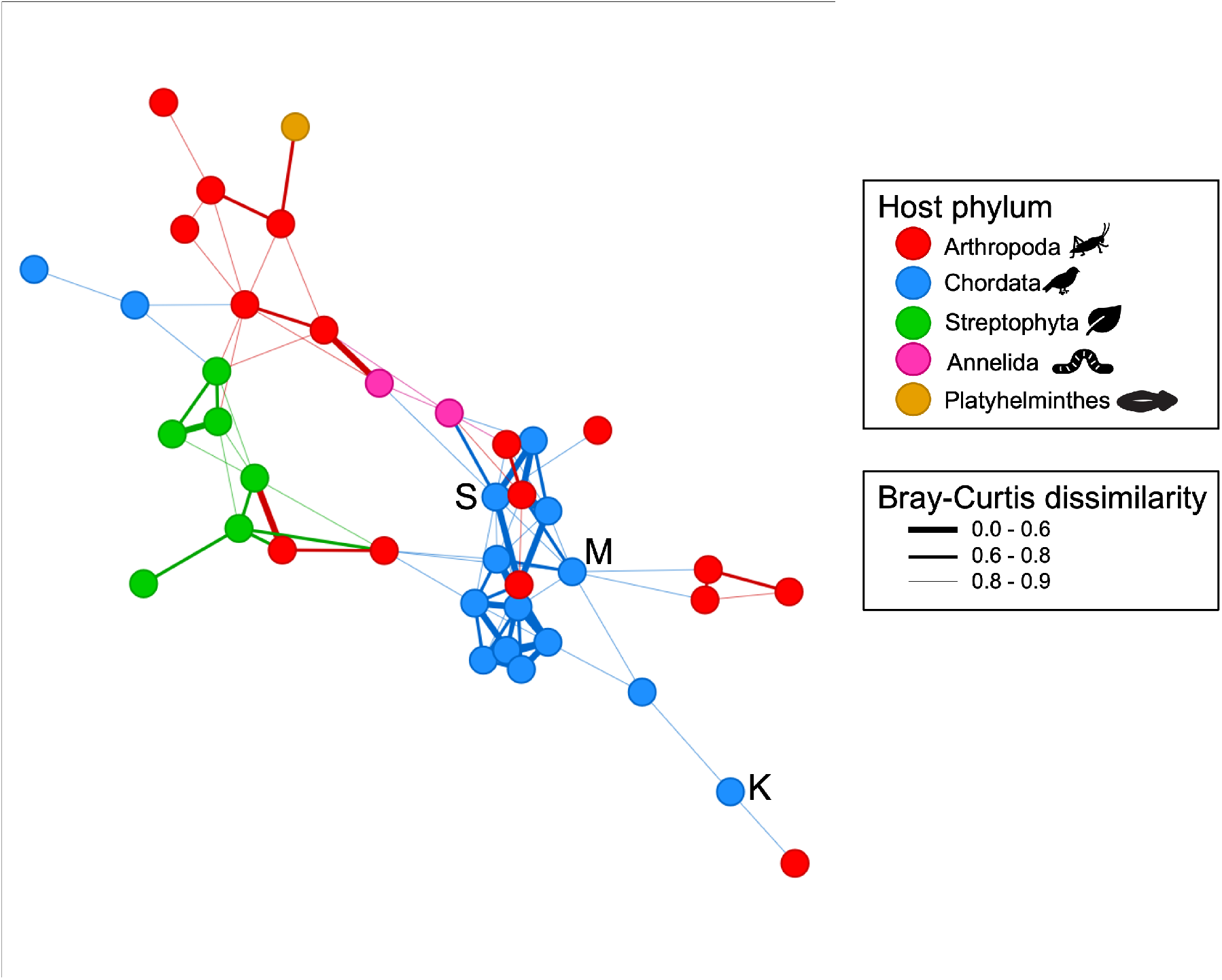
Host community network (unipartite) displayed using the Fruchterman-Reingold layout. Nodes are connected to other nodes if they have a dissimilarity value of less than 0.9. The thickness of the line shows the level of dissimilarity. The Bray-Curtis dissimilarity statistic ranges from 0 to 1, with 0 meaning two hosts have identical viromes at the viral family level, and 1 meaning two hosts have no viral families in common. The letters refer to species of interest; K=kākāpō, M=mohua, S=Te Kakahu skink.

In comparison to a null network, the transitivity ratio (i.e., the probability that the adjacent nodes are connected, expressed as a ratio comparing the null and host community network values) was high (3.1), while the path length ratio (the smallest number of links between each node) was similar (1.2). This signifies a small world network: both highly clustered and highly connected. Most nodes are a small number of links away from every other node, enabling viruses to easily move between species (*17*). Our analysis also revealed that if a virus were to move through the network, it would first move within a host phylum, then most likely from plants to arthropods, and to/from arthropods and chordates. Strikingly, direct links between plants and chordates were far less common, which may reflect the relatively small number of chordates on the island that eat plants, compared to those that eat invertebrates (table S1).

### Host ecology affects viral diversity at smaller scales

To examine the diversity of virus species within and across host phyla we conducted phylogenetic analysis for one virus family per module: *Parvoviridae* (module 1), *Caulimoviridae* (module 2), *Fiersviridae* (module 3) and *Caliciviridae* (module 4) (Fig. 6, fig. S2–S5, table S6), chosen for their high viral diversity across multiple host species. In general, there was a high degree of virus clustering by host phyla, with each module containing a high richness and abundance of virus species from their predominant host. However, there was also evidence of host ecology (i.e., predator-prey interactions) influencing virus diversity, with closely related/identical viruses found in distantly related hosts. For example, we identified three viruses in the kākā (parrot) library from the plant virus family *Caulimoviridae* that clustered close to members of the viral genus *Badnavirus* found in the plants sampled here (Fig. 6). This pattern is indicative of a food web interaction in which the parrot consumed the plants. Similarly, we found near identical (>99% at the amino acid level) members of the *Caliciviridae* in a miromiro/tomtit (passerine, *Petroica macrocephala*) and slater/woodlouse (arthropod, *Isopoda* sp.) library, strongly indicating a food web interaction (fig. S5). As the arthropod is a detritivore and the bird predominately insectivorous, the interaction could have occurred in either or both directions.

**Fig. 6.**
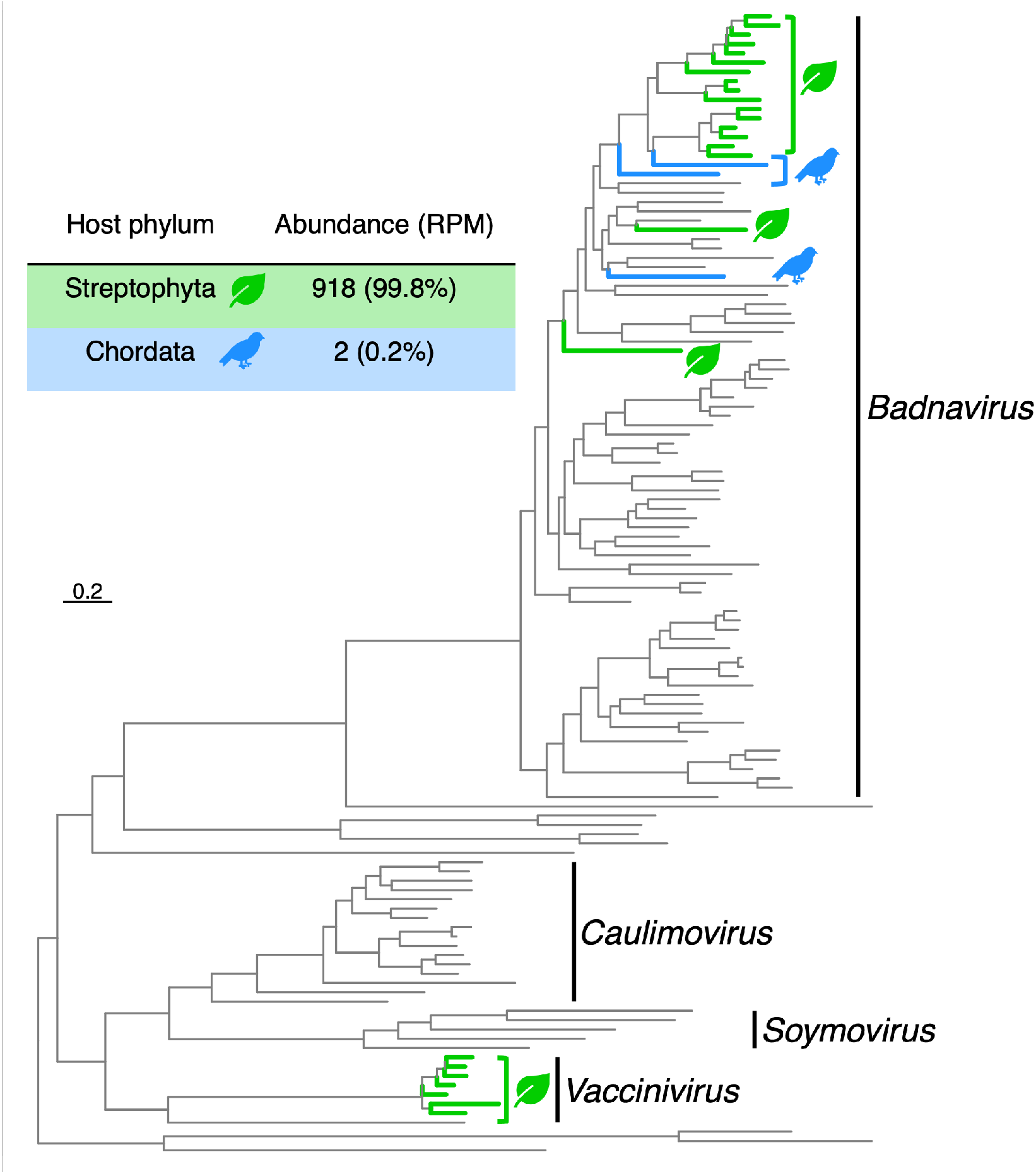
Virus diversity at the species level. Phylogeny of the *Caulimoviridae* (module 2). The colours and symbols correspond to host phyla, green leaf = Streptophyta, blue bird = Chordata. All unmarked viruses have plant hosts. The abundance of viruses in each phylum is shown in the table inset, expressed as reads per million (RPM). Branches are scaled according to the number of amino acid substitutions per site, shown in the scale bar. The tree is midpoint rooted for display purposes only. Detailed individual phylogenies, sequence alignments, and information including the genes used, alignment length, percentage identity and number of sequences can be found in tables S5-S6 and fig. S2-S5.

## Discussion

We used a metatranscriptomic approach to reveal the drivers of virome structure in a relatively isolated island community. We achieved this by creating the host-virome network (bipartite) based on the presence/absence of each virus family in each host library, and the host community network (unipartite), based on the level of virome similarity between hosts.

This analysis revealed a strong effect of host phylogeny on the host-virome network, with more viral families shared within than between host phyla. Such a clear effect provides strong evidence that host phylogenetic distance is a key constraint on the cross-species transmission of viruses, and hence on disease emergence. Similar trends have been observed at smaller scales (i.e., with specific viral families and hosts), with cross-species transmission decreasing in frequency with increasing host phylogenetic distance (*18, 19*). Host phylogeny has also been shown to be an important factor in explaining the diversity of bacteria and eukaryotic parasites (*20, 21*), indicating that it has wide ranging impacts on all infectious agents within an organism. The strength of this host phylogenetic trend is surprising given that the viromes characterised included all viruses, not just those infecting the host in question. There were necessarily differences in sampling method between chordates (cloacal swab), invertebrates (tissue) and plants (tissue) which, while unavoidable, could have resulted in artificial similarities between similar sample types. However, cloacal swabs should have biased the results toward food web interactions since they represent a sample of the host and the digestive tract, which often contain diet-associated viruses (*22*).

That viruses generally cluster by host phylogeny means that virus-host co-divergence will shape virome structure on deep evolutionary scales. Phylogenetic barriers may prevent frequent cross-species virus transmission between phyla because this can only occur between species with similar virus-cell receptor interactions. This results in a mixture of host-virus codivergence at deeper taxonomic levels and cross-species transmission at shallower taxonomic boundaries, with the combination of both processes meaning that more closely related species have generally similar viromes. This is in accord with most comparative studies that have revealed a relatively weak impact of host-virus codivergence (*23, 24*). Indeed, our phylogenetic analysis at the level of virus species provided direct evidence for cross-species transmission with, for example, a cluster of closely related caliciviruses in both the fantail (*Rhipidura fuliginosa*, passerine) and the tawaki (*Eudyptes pachyrhynchus*, penguin) (fig. S5).

A striking result was the limited impact of host diet on the host-virome network within Chordata, explaining only a small amount of variation. This suggests that predator-prey interactions are not associated with high levels of cross-species viral transmission, likely due to the constraints associated with phylogenetic barriers. Aside from the human gut (*25*), there is limited evidence for host diet impacting host viromes. However, other aspects of host ecology such as location, age and behaviour (within a narrower host range than our study) all influence host viromes, although to a lesser extent than phylogeny (*4, 26*). Despite the overall weak influence of host ecology on viral diversity, we found clear examples of host predator-prey interactions at the virus species level, showing that viruses are moved between species via these processes. Overall, our results suggest that virus traffic from predator-prey interactions is only transient (i.e., the virus is only present for a short time in the predator’s digestive system) and unlikely to result in productive infections.

Both the host-virome network (bipartite) and host community network (unipartite) had similar structures to other ecological networks, suggesting they have similar constraints. Our host-virome network followed a truncated power-law distribution that had lower cut-off values than expected by chance. This suggests that the structure is influenced by assembly mechanisms (i.e., ecological or phylogenetic traits of the host or virus that alter how the network is constructed). Power-law distributions occur when nodes are likely to get more links the more they have already, such that the ‘rich get richer’ (*27*). Accordingly, hosts that already have a high diversity of viruses are more likely to gain new viruses than hosts with a low diversity of viruses. Similarly, viruses with many hosts are more likely to gain more hosts, which may inform zoonotic risk assessments (*28*): rather than assigning the highest risk score to viruses that are similar to those that have already emerged, it may be of greater utility to give preference to the most generalist viruses. This power-law distribution is truncated when the underlying model no longer accurately predicts the distribution beyond a certain cut-off, effectively preventing the rich from getting richer beyond that point. The higher this cut-off, the greater the number of highly connected hosts and viruses (*29*). In our models, this cut-off value was higher than for the null network, indicating that the host-virome network contains more highly connected hosts and viruses than expected by chance. Truncated power-law distributions are often found in mutualistic networks (e.g. plant-pollinators), with the truncation thought to be due to ‘forbidden’ – physically impossible – links (*30*). In our network, forbidden links could be due to phylogenetic barriers preventing cross-species transmission.

The host community network (unipartite) was a ‘small world’ network. Although clear clusters emerge, most nodes (hosts) can be reached via a small number of connections to other nodes (*31*). This is common in ecological networks, with most species only two links apart on average in complex food webs (*32*). The combination of strong modularity in our host virome network and high connectivity in our host community network implies that a pathogenic virus could rapidly move through the network, particularly once the virus crossed from one module to another. The host community network showed an especially strong clustering within chordates, with a high degree of similarity between hosts. This cluster included endangered species such as the mohua and Te Kakahu skink, suggesting that on Pukenui/Anchor Island these species are vulnerable to disease emergence from other chordates within that cluster. In contrast, the kākāpō was less closely connected to this cluster, suggesting it may be less vulnerable.

Our study shows that the phylogenetic relatedness of hosts is a strong driver of viral diversity in ecological communities. Phylogenetic barriers between distantly related hosts likely prevent frequent virus movement despite exposure events via predator-prey interactions. The ecological community is highly connected, which carries risks for disease emergence in vulnerable species. Our study sheds light on the processes that dictate viral movement through ecosystems. Further research is needed on the implications of disruption to these networks on disease emergence.

## Acknowledgments

We thank Andrew Digby, Jodie Crane, Brodie Philp, Dave Johnson and Chloe Corne for logistical assistance, and Brodie Philp, Bex Jackson and Nigel French for assistance with sample collection. We thank the Molecular Epidemiology and Public Health Laboratory at Massey University New Zealand for use of their laboratory space, and Kristene Gedye for advice and assistance. This research utilised the high-performance computing service, Artemis, provided by the Sydney Informatics Hub, Core Research Facility, University of Sydney.

## Funding

Australian Research Council Australian Laureate Fellowship FL170100022 (ECH). FRAGCLIM ERC Consolidator Grant under the European Union’s Horizon 2020 (Grant Agreement Number 726176), and “Laboratoires d’Excellences (LABEX)” TULIP (ANR-10-LABX-41) (JMM)

## Author contributions

Conceptualization: RKF, MWT, KRT, ECH

Methodology: RKF, MM, MWT, KEC, SHA, CMM, KRT, ECH

Investigation: RKF, MM, KEC, SHA, TCG, CMM, CGM, KRT

Formal analysis: RKF, JMM

Visualization: RKF

Funding acquisition: ECH, JMM

Supervision: ECH

Writing – original draft: RKF

Writing – review & editing: RKF, MM, SHA, KEC, TCG, CMM, CGM, MWT, JMM, KRT, ECH

## Competing interests

Authors declare that they have no competing interests.

## Data and materials availability

The operational taxonomic unit table used in analyses is provided in Data S1. The non-host sequence data generated in this study has been deposited in the Sequence Read Archive (SRA) under the accession numbers SAMN30927701-49. Virus consensus sequences have been submitted to NCBI/GenBank and assigned accession numbers XXXX-YYYY.

## Supplementary Materials

Materials and Methods

Figs. S1 to S5

Tables S1 to S6

Data S1 (separate file)

References (*33-56*)

## Notes

### Competing Interest Statement

The authors have declared no competing interest.

