## Supplementary Information for "Host Phylogeny Shapes Viral Transmission Networks in an Island Ecosystem"

### Host Phylogeny Shapes Virus Evolution in an Island Ecosystem

**This PDF file includes:**

Materials and Methods
Figs. S1 to S5
Tables S1 to S6
Caption for Data S1

**Other Supplementary Materials for this manuscript include the following:**

Data S1 [OTU_table_raw_abundances.xlsx]

### Materials and Methods

Study location

Pukenui/Anchor Island is a small island (11.4 $km^2$), located in Dusky Sound, Fiordland, New Zealand (fig. S1). The island is part of the largely uninhabited Fiordland National Park (over 12,000$km^2$) on the south-west coast of the South Island and is over 80 km to the nearest township by air. Following the eradication of invasive mammals in the early 2000s, the island became key habitat for endangered native species, including the kākāpō (*Strigops habroptila*). The island is also an important habitat for seabirds that nest in the forest, including the tawaki/Fiordland crested penguin (*Eudyptes pachyrhynchus*). The temperate rainforest consists predominately of beech and podocarp (conifer) trees, with an understory/forest floor including shrubs, vines and mosses. To our knowledge, the only non-native permanent inhabitant of the island is the invasive German wasp (*Vespula germanica*).

Sample collection

This research was conducted under a Department of Conservation Wildlife Act Authority Authorisation number 86173-FAU, Authority for research and/or collection of material on public conservation land Authorisation number 86172-RES and had ethics approval from the University of Auckland reference number 002198.

Fieldwork was undertaken on Anchor Island, Fiordland, New Zealand between the 17$^{th}$ of February and 14$^{th}$ of March 2021. The 18 bird and 1 skink species were caught using four different methods, depending on the species in question. Small, flighted birds were caught using low canopy mist-netting, while larger flighted birds were caught with high canopy mist-nets. Bird calls were used to attract the birds to the area and into the nets. Non-flying birds were caught by hand or hand-net. Skinks were caught using gee-minnow traps. Once caught, the animals were weighed and a cloacal swab was taken, using a sterile nylon flocked swab FLOQswab$^{TM}$ (Copan), either mini-tip or regular-tip depending on the size of the animal. The entire tip of the swab was inserted into the cloaca and swabbed with two to four circular motions while applying gentle pressure against the mucosal surfaces. The swab was then cut using scissors sterilised with 70% alcohol and placed into a tube with 1 ml of RNAlater. Samples were kept at -20°C for the duration of the fieldwork, then frozen at -80°C.

Leaves were collected from each plant species by cutting the stem with sterile scissors and placing the leaves into a sterile collection bag (1 bag per individual plant). At the fieldwork base, a leaf from each individual plant was chopped into approximately 5 mm x 5 mm pieces and placed into 1 ml of RNAlater. The total volume of the solution and plant material was no more than 1.3 ml, to ensure preservation of all the RNA. To ensure the RNAlater permeated into the tissue, samples were left at 4°C for approximately 12 hours before being transferred to -20°C. Samples were kept at -20°C for the duration of the fieldwork, then frozen at -80°C.

Invertebrates were collected in two ways – by manual search, and by extraction from soil. At five sites on Anchor Island, the area within a 5 x 5 metre square was intensively searched for invertebrates (on vegetation, under logs, under bark etc.). When an invertebrate was found it was placed alive into a sterile pottle with damp moss/leaf litter from the site. At the same site, a sterilised spade was used to cut a soil core approximately 2 litres in volume. The core was placed into a sterilised 2 litre container. The invertebrates and soil cores were kept at 4°C for the duration of the fieldwork. They were then transferred to the Ecology Department at Massey University, New Zealand. The invertebrates collected by hand were examined live under a dissecting microscope using sterile tools and identified to the lowest classification level possible (highest = order level, lowest = species level). Invertebrates from the soil cores were extracted using Berlese funnels into RNAlater and identified in the same way. Once identified, the invertebrates were individually stored at -80°C.

RNA extraction – cloacal swabs

RNA was extracted using the RNeasy plus mini extraction kit (Qiagen) and QIAshredders (Qiagen). The tube containing the swab in RNAlater was thawed and the swab removed from the tube using sterile forceps and placed in 600 µl of extraction buffer. The swab and buffer were vortexed for two minutes at maximum speed. The swab and buffer were then placed into a QIAshredder and centrifuged for five minutes at maximum speed. The flowthrough was retained (avoiding the cell debris pellet) and used in the extraction following the standard protocol in the kit. The RNA was eluted into 50 µl of sterile water. Extractions were pooled by host species for sequencing. 25 µl of each extraction were used in each pool, and this was concentrated using the NucleoSpin RNA Clean-up XS, Micro kit for RNA clean up and concentration (Macherey-Nagel). The concentrated RNA was eluted into 20 µl of sterile water.

RNA extraction – Plant material

RNA was extracted using the RNeasy plant mini extraction kit (Qiagen). The plant material in RNAlater was thawed just enough to remove approximately 20–30 mg. This was placed into a tube with a sterile stainless steel 6 mm bead. The tube, sample, bead and adapter set were then cooled at -80°C for 30 minutes. After cooling, the plant tissue was disrupted by beating using the TissueLyser II (Qiagen) at 30Hz for two minutes. The kit protocol was then followed for the remainder of the extraction. The RNA was eluted in 50 µl of sterile water. Extractions were pooled and concentrated as described above. Before concentrating the pooled RNA was treated with DNase, using the rDNase Set (Macherey-Nagel).

RNA extraction – Invertebrates

RNA was extracted using the RNeasy plus mini extraction kit (Qiagen). For small invertebrates (<30 mg), the whole body was used in the extraction. For larger invertebrates a 30 mg piece of the abdomen was used. The frozen tissue was placed into a tube with a sterile stainless steel 6 mm bead and 300-600 µl of buffer was added, depending on the amount of material. The tissue was disrupted by beating using the TissueLyser II (Qiagen) at 30 Hz for 4 minutes. The kit protocol was then followed for the remainder of the extraction. The RNA was eluted in 50 µl of sterile water. Extractions were pooled and concentrated as described above.

Total RNA Sequencing

cDNA libraries were prepared using the Stranded Total RNA Prep with Ribo-Zero Plus (Illumina) for cloacal swabs and invertebrates, and the TruSeq Stranded Total RNA with Ribo-Zero Plant (Illumina) for plants. Libraries were sequenced on the Illumina Novaseq platform, with invertebrates, plants and vertebrates sequenced entirely independently (i.e. on different lanes and sequencing runs). One blank negative control library (i.e., a sterile water and reagent mix) was sequenced with each sequencing run (one each for vertebrates, invertebrates and plants).

Quality control, assembly and virus identification

Using Trimmomatic (0.38) (*33*), adapters and bases below a quality of five were trimmed, using a sliding window approach with a window size of four. Bases were cut if below a quality score of three at the beginning and end of the reads. Using bbduk in BBtools (bbmap

37.98) (*34*), sequences less than 100 nucleotides in length or below an average quality of ten were removed.

Reads were *de novo* assembled using Megahit (1.2.9) (*35*). Viruses were identified by comparing the assembled contigs to the NCBI nucleotide database (nt) and non-redundant protein database (nr) using Blastn (blast+ 2.1.2) (*36*) and Diamond Blastx (Diamond 2.0.9) (*37*). Contigs were retained that had hits to viruses and an open reading frame greater than 300 nucleotides (for nr hits). Sequence similarity cut-off values of 1E-5 and 1E-10 were used for the nt and nr databases, respectively, to prevent false-positives. Virus abundance was estimated using Bowtie2 (2.2.5) (*38*). Viruses that met the following conditions: (i) sequenced on the same lane, (ii) the total read count was < 0.1% of the read count in the other library, and (iii) were >99% identical at the nucleic acid level were assumed to be contamination due to index-hopping from another library and removed. Any virus found in the blank negative control libraries was assumed to have resulted from contamination and similarly removed from all libraries and analyses.

<u>Ecological analysis</u>

All analyses were conducted in R (4.0.5) (*39*). Viruses were grouped into viral families, as classified by the International Committee on Taxonomy of Viruses (ICTV) or National Center for Biotechnology Information (NCBI). Viruses not classified to family level were included provided they had an order-level classification (e.g., unclassified *Picornavirales* were included as a 'family'). An OTU (Operational Taxonomic Unit) table was created using viral abundance expressed as the number of reads per million (RPM, i.e., the number of reads from the virus family divided by the total number of reads in the library, multiplied by one million). Using this table, beta diversity (i.e., shared diversity across host phyla) was visualised at the virus family level using non-metric multidimensional scaling with a Bray–Curtis dissimilarity matrix, presented as an ordination plot using the R package phyloseq (*40*) and scatterplot3d (*41*). Bray–Curtis dissimilarity is a statistic ranging from 0 to 1 that reflects the dissimilarity of communities between libraries, with 0 meaning both libraries have an identical community, and 1 meaning the libraries have no virus families in common. Pairwise permutational analyses of variance (PERMANOVA, adonis2 in the vegan R package (*42*), and pairwise adonis test from pairwiseAdonis (*43*)) were used to test for differences in the community based on host phylogenetic groupings and host ecology, using the Bray-Curtis dissimilarity matrix and an alpha of 0.05 following a Bonferroni correction. Where multiple terms were used, the marginal effects of each term were tested using by="margin" in adonis2.

A bipartite network was constructed using igraph (*44*) and visualised using Visnetwork (*45*), based on the presence/absence of each virus family in each host library. Two sets of nodes were defined: virus families and host species. One link in the network corresponded to a virus node inhabiting a host node whenever the virus family was found in the host library. This host-virome network comprised 49 host libraries, 112 virus families and 926 interactions between these two node sets. Modules (i.e., groups of nodes with more links among them than to the rest of the network, also named communities) were identified to infer the community structure in the host-virome network. To do this we used the DIRTLPAwb+ community detection algorithm (*46*) in the bipartite package (*47*) that identifies partitions with high modularity scores by maximising weighted modularity, weighted using log abundance (not RPM). DIRTLPAwb+ employs multiple iterations of the LPBwb+ algorithm, based on Barber's modularity (*48*):

$$Q = \frac{1}{2m} \sum (A_{ij} - P_{ij}) \delta(g_i, g_j)$$

In which $Q$ is the modularity score, $m$ is the number of links in the network, $g_i$ and $g_j$ are the assigned modules for nodes i and j. $A_{ij} = 1$ if a link exists between nodes i and j, or 0 if no link. $P_{ij}$ is the probability that a link exists between nodes i and j based on a null model. $\delta(g_i, g_j) = 1$ if the modules are the same, and 0 if different (*46, 48*).

Pairwise adonis tests were used to test for differences in community composition between modules, using module as an independent factor and an alpha of 0.05 following Bonferroni correction.

We evaluated the robustness of the modules identified by rarefying to the lowest sequencing depth following the methods used by Lurgi *et al.* (*49*). We performed pairwise adonis tests on 100 rarefied data sets of the original data using the modules detected in our host-virome network as an independent variable. We rarefied the data to the size of the smallest library (630 contigs) using the rrarefy function of the vegan package (*42*). This was repeated 100 times independently to obtain 100 different rarefied data sets, using the replicate function in base-R. We then analysed each of these rarefied data sets with the pairwise adonis test and averaged the result across the 100 rarefied data sets. In this way, we tested whether the communities identified by the modularity analysis remained consistently significantly different when rarefied down to the lowest sequencing depth.

Using the modules obtained from the community detection algorithm, we evaluated the roles of individual species in the network by analysing the degree distribution (the distribution of the number of links from each node). To assess the distribution of links between nodes in the host-virome network we also calculated the cumulative degree distribution and fitted a truncated power law, using nls and the Kolmogorov-Smirnov test to evaluate whether it fitted the power law distribution. The truncated power law distribution was as follows:

$$Pc(k) = k^{i}\, k^{-(\frac{k}{z})}$$

Where $k$ is the cumulative degree distribution, $i$ is the power-law decay exponent and $k^{-(\frac{k}{z})}$ is the exponential cut-off for the truncation, where $z$ is the cut-off value beyond which the power-law distribution no longer fits.

We then compared this distribution to a null model of a random bipartite graph created using the Erdos-Renyi model, with the same number of host and virus nodes and interactions, but with the interactions randomized.

To examine the strength of connections between hosts created by shared viromes, we created a unipartite network (i.e., links between hosts with shared virus families) based on the Bray-Curtis dissimilarity matrix, using the phyloseq and igraph R packages. Given the Bray-Curtis statistic ranges from 0 to 1, a cut-off was required to determine how similar two communities need to be to link them in the network. We chose a cut-off of 0.9, which is the lowest that still creates a cohesive network (i.e., no isolated groups of nodes). However, this cut-off does create nine singletons (nodes with no connections), which were removed. The network (referred to as the 'host community' network) comprised 40 hosts and 99 interactions. The 'small world' properties of this network were examined using the distance_table functions in igraph, including calculating the shortest paths (the smallest number of links between each node) and average shortest path length (the mean of all the shortest paths). We used the smallworldness function in qgraph (*50*), which uses the transitivity (the probability that the adjacent nodes of a vertex are connected, using the definition and formula developed by Barrat (*51*)) and average shortest path length to compare the network to randomly generated networks.

<u>Phylogenetic analysis</u>

The patterns of virus diversity within viral families were visualised using phylogenetic trees with one viral family per module examined in detail, out of a total of 112 viral families. Amino acid sequences were aligned using MAFFT (7.402) (*52*) employing the L-INS-i algorithm and trimmed with a gap threshold of 0.9 and at least 20% of the sequence conserved using TrimAl (1.4.1) (*53*). Individual maximum likelihood phylogenetic trees for each virus family were estimated using IQ-TREE (1.6.12) (*54*), with the best-fit substitution model determined by the program and node robustness assessed by employing the approximate likelihood ratio test with 1000 replicates. Phylogenetic trees were visualised using APE (5.4) (*55*) and ggtree (2.4.1) (*56*) in R.

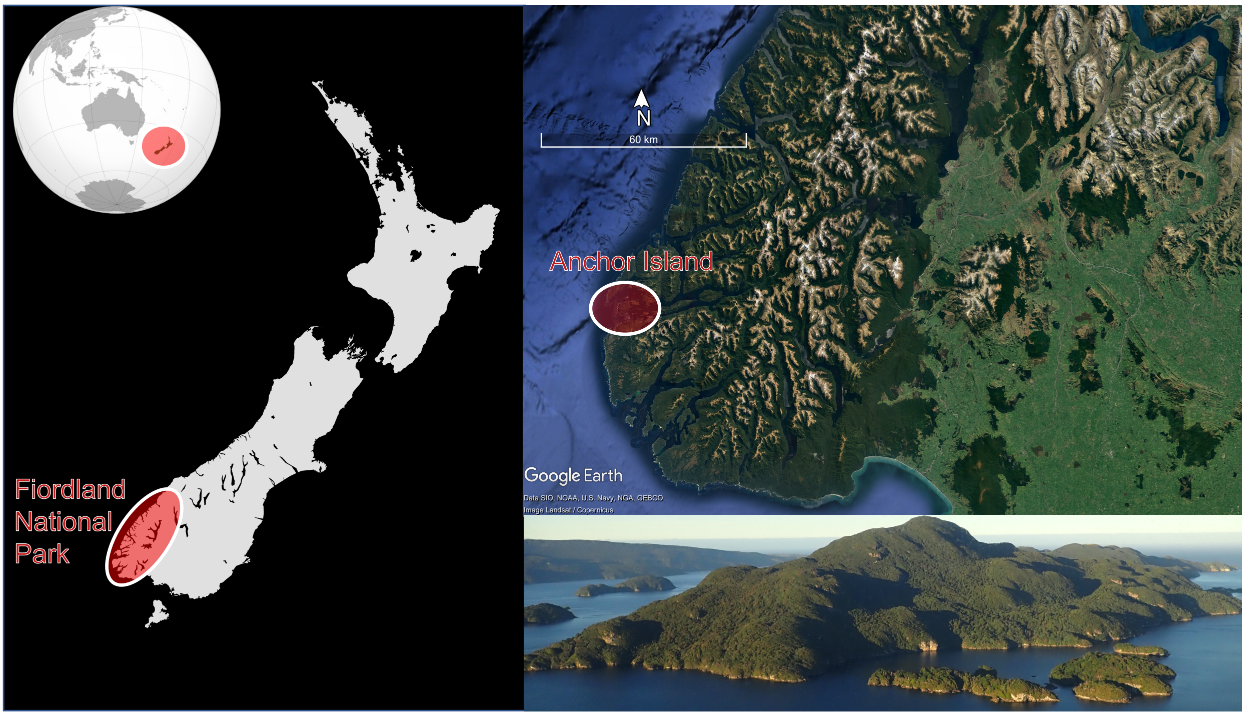


**Fig. S1.** Map showing the location of Anchor Island, New Zealand. Left – Anchor Island is in the Fiordland National Park in the south-west corner of the South Island, New Zealand. Top right – The island is at the seaward end of Dusky Sound. Bottom right – The island is steep and entirely covered in native vegetation.

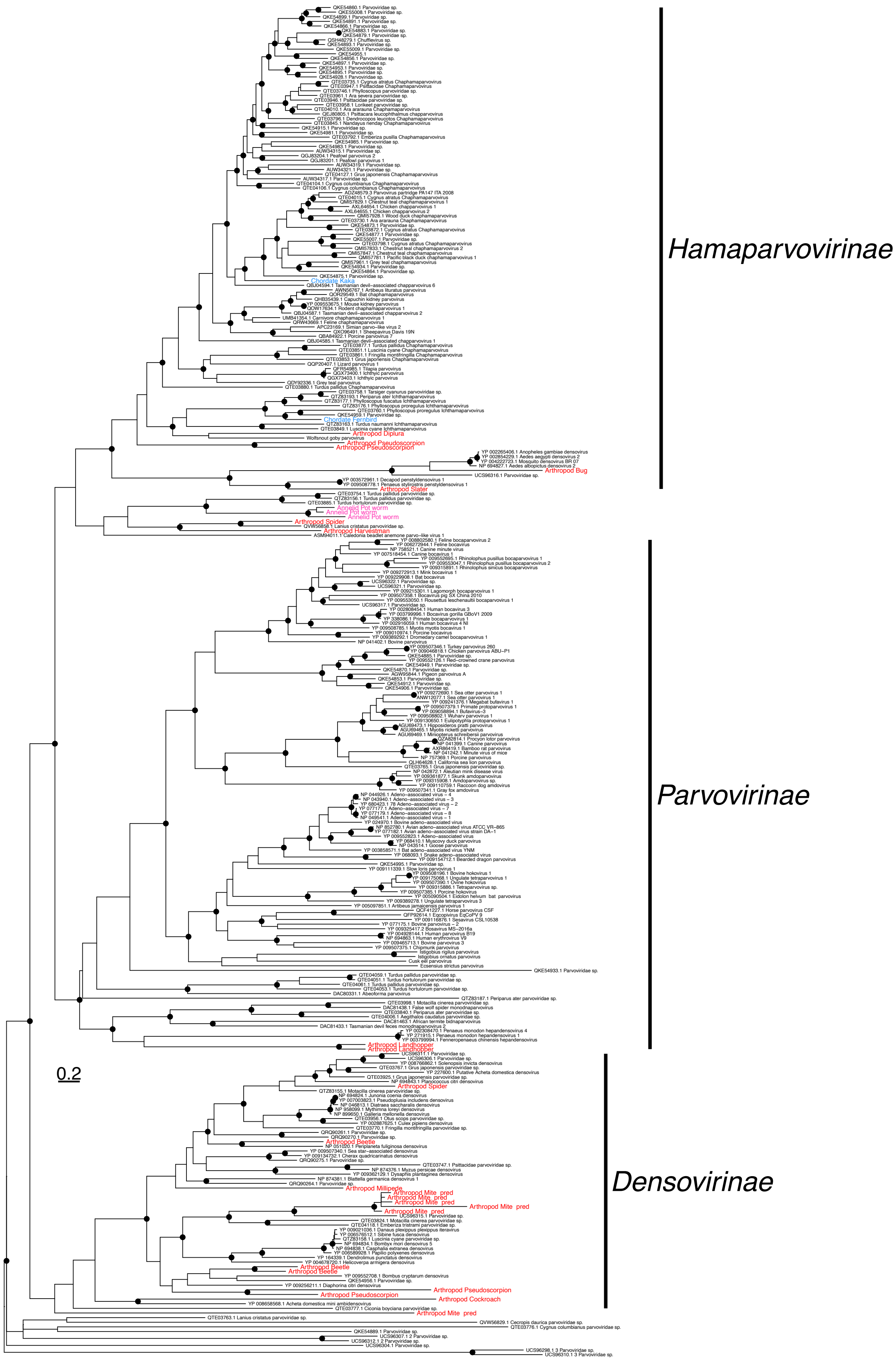
Hamaparvovirinae
Parvovirinae
Densovirinae
0.2

**Fig. S2**. Phylogenetic trees for the *Parvoviridae* based on the non-structural protein 1 gene (alignment length 584 amino acids). The colours correspond to host phyla, blue = Chordata, red = Arthropoda, pink = Annelida. Viruses from this study are labelled with their phylum and host common name (see Table S1 for details about each host). Branches are scaled according to the number of amino acid substitutions per site, shown in the scale bar. Black circles on nodes show bootstrap support values of more than 90%. The tree is midpoint rooted for display purposes only.

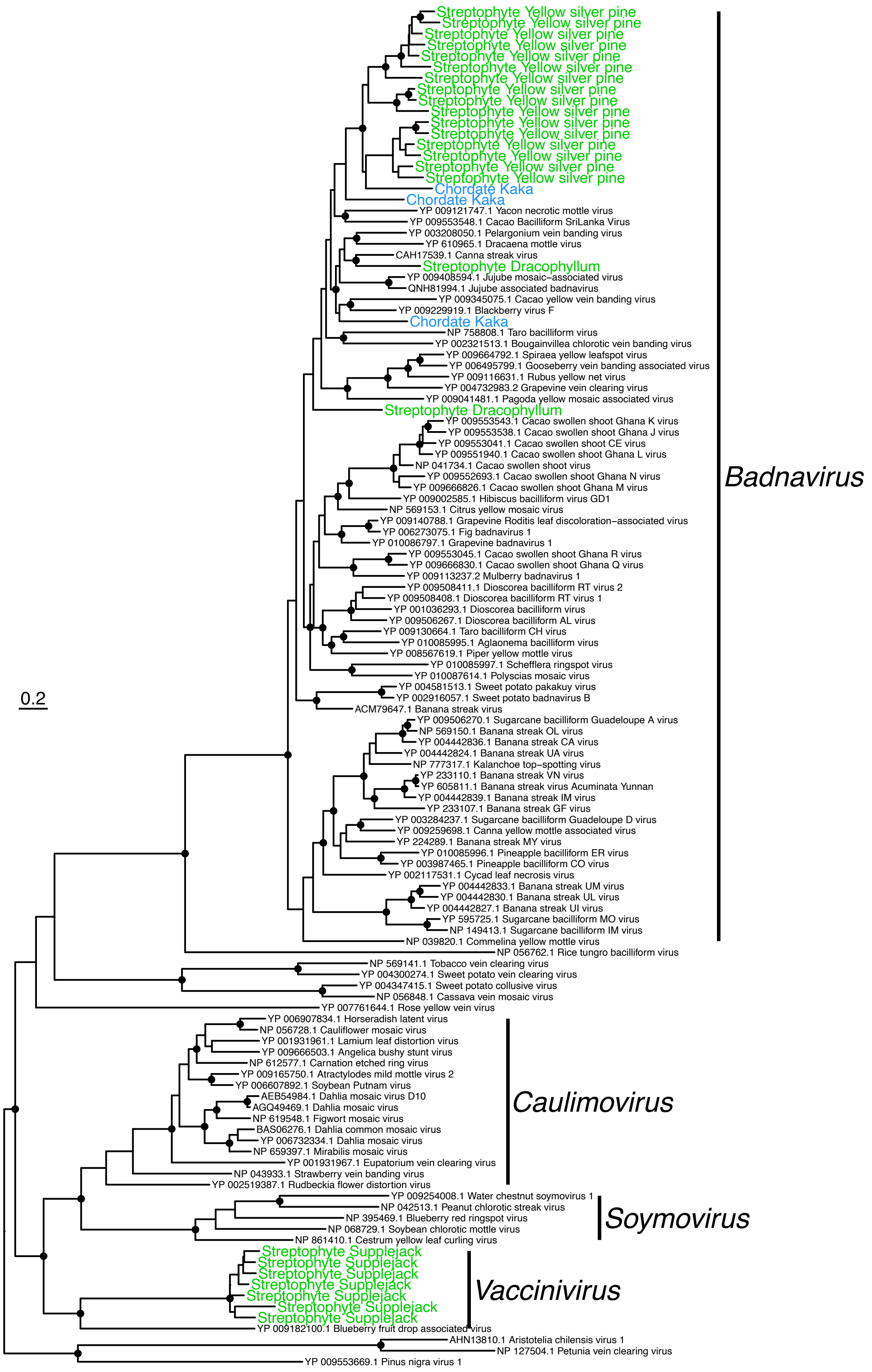
0.2
Badnavirus
Caulimovirus
Soymovirus
Vaccinivirus
Streptophyte Yellow silver pine
Chordate Kaka
YP 009121747.1 Yacon necrotic mottle virus
YP 009553548.1 Cacao Bacilliform SriLanka Virus
YP 003208050.1 Pelargonium vein banding virus
YP 610965.1 Dracaena mottle virus
CAH17539.1 Canna streak virus
Streptophyte Dracophyllum
YP 009408594.1 Jujube mosaic–associated virus
QNH81994.1 Jujube associated badnavirus
YP 009345075.1 Cacao yellow vein banding virus
YP 009229919.1 Blackberry virus F
NP 758808.1 Taro bacilliform virus
YP 002321513.1 Bougainvillea chlorotic vein banding virus
YP 009664792.1 Spiraea yellow leafspot virus
YP 006495799.1 Gooseberry vein banding associated virus
YP 009116631.1 Rubus yellow net virus
YP 004732983.2 Grapevine vein clearing virus
YP 009041481.1 Pagoda yellow mosaic associated virus
YP 009553543.1 Cacao swollen shoot Ghana K virus
YP 009553538.1 Cacao swollen shoot Ghana J virus
YP 009553041.1 Cacao swollen shoot CE virus
YP 009551940.1 Cacao swollen shoot Ghana L virus
NP 041734.1 Cacao swollen shoot virus
YP 009552693.1 Cacao swollen shoot Ghana N virus
YP 009666826.1 Cacao swollen shoot Ghana M virus
YP 009002585.1 Hibiscus bacilliform virus GD1
NP 569153.1 Citrus yellow mosaic virus
YP 009140788.1 Grapevine Roditis leaf discoloration–associated virus
YP 006273075.1 Fig badnavirus 1
YP 010086797.1 Grapevine badnavirus 1
YP 009553045.1 Cacao swollen shoot Ghana R virus
YP 009666830.1 Cacao swollen shoot Ghana Q virus
YP 009113237.2 Mulberry badnavirus 1
YP 009508411.1 Dioscorea bacilliform RT virus 2
YP 009508408.1 Dioscorea bacilliform RT virus 1
YP 001036293.1 Dioscorea bacilliform virus
YP 009506267.1 Dioscorea bacilliform AL virus
YP 009130664.1 Taro bacilliform CH virus
YP 010085995.1 Aglaonema bacilliform virus
YP 008567619.1 Piper yellow mottle virus
YP 010085997.1 Schefflera ringspot virus
YP 010087614.1 Polyscias mosaic virus
YP 004581513.1 Sweet potato pakakuy virus
YP 002916057.1 Sweet potato badnavirus B
ACM79647.1 Banana streak virus
YP 009506270.1 Sugarcane bacilliform Guadeloupe A virus
NP 569150.1 Banana streak OL virus
YP 004442836.1 Banana streak CA virus
YP 004442824.1 Banana streak UA virus
NP 777317.1 Kalanchoe top–spotting virus
YP 233110.1 Banana streak VN virus
YP 605811.1 Banana streak virus Acuminata Yunnan
YP 004442839.1 Banana streak IM virus
YP 233107.1 Banana streak GF virus
YP 003284237.1 Sugarcane bacilliform Guadeloupe D virus
YP 009259698.1 Canna yellow mottle associated virus
YP 224289.1 Banana streak MY virus
YP 010085996.1 Pineapple bacilliform ER virus
YP 003987465.1 Pineapple bacilliform CO virus
YP 002117531.1 Cycad leaf necrosis virus
YP 004442833.1 Banana streak UM virus
YP 004442830.1 Banana streak UL virus
YP 004442827.1 Banana streak UI virus
YP 595725.1 Sugarcane bacilliform MO virus
NP 149413.1 Sugarcane bacilliform IM virus
NP 039820.1 Commelina yellow mottle virus
NP 056762.1 Rice tungro bacilliform virus
NP 569141.1 Tobacco vein clearing virus
YP 004300274.1 Sweet potato vein clearing virus
YP 004347415.1 Sweet potato collusive virus
NP 056848.1 Cassava vein mosaic virus
YP 007761644.1 Rose yellow vein virus
YP 006907834.1 Horseradish latent virus
NP 056728.1 Cauliflower mosaic virus
YP 001931961.1 Lamium leaf distortion virus
YP 009666503.1 Angelica bushy stunt virus
NP 612577.1 Carnation etched ring virus
YP 009165750.1 Atractylodes mild mottle virus 2
YP 006607892.1 Soybean Putnam virus
AEB54984.1 Dahlia mosaic virus D10
AGQ49469.1 Dahlia mosaic virus
NP 619548.1 Figwort mosaic virus
BAS06276.1 Dahlia common mosaic virus
YP 006732334.1 Dahlia mosaic virus
NP 659397.1 Mirabilis mosaic virus
YP 001931967.1 Eupatorium vein clearing virus
NP 043933.1 Strawberry vein banding virus
YP 002519387.1 Rudbeckia flower distortion virus
YP 009254008.1 Water chestnut soymovirus 1
NP 042513.1 Peanut chlorotic streak virus
NP 395469.1 Blueberry red ringspot virus
NP 068729.1 Soybean chlorotic mottle virus
NP 861410.1 Cestrum yellow leaf curling virus
Streptophyte Supplejack
YP 009182100.1 Blueberry fruit drop associated virus
AHN13810.1 Aristotelia chilensis virus 1
NP 127504.1 Petunia vein clearing virus
YP 009553669.1 Pinus nigra virus 1

**Fig. S3**. Phylogenetic tree of the *Caulimoviridae* based on the polyprotein gene (alignment length 938 amino acids). The colours correspond to host phyla, green = Streptophyta, blue = Chordata. Viruses from this study are labelled with their phylum and host common name (see Table S1 for details about each host). Branches are scaled according to the number of amino acid substitutions per site, shown in the scale bar. Black circles on nodes show bootstrap support values of more than 90%. The tree is midpoint rooted for display purposes only.

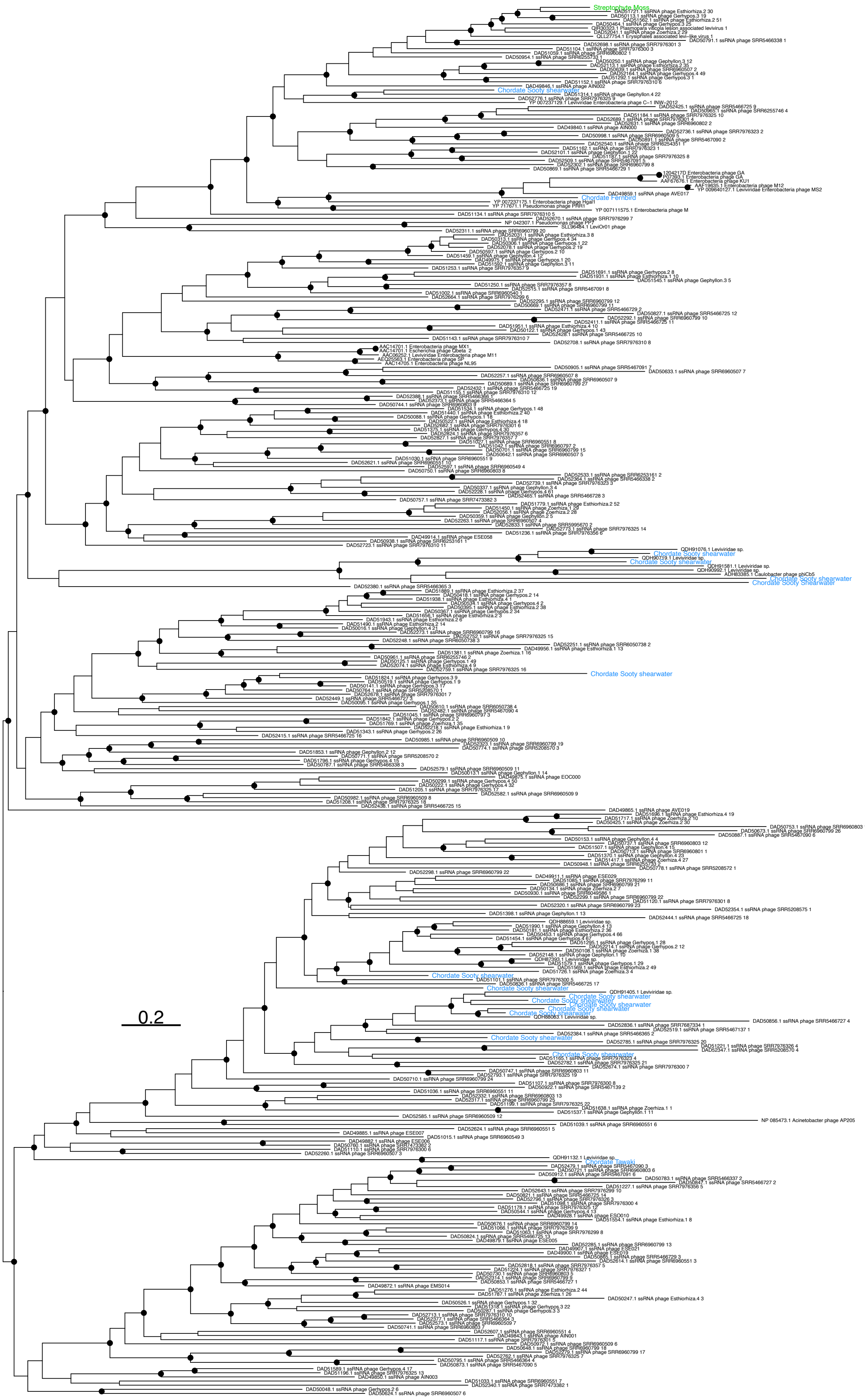
0.2

**Fig. S4**. Phylogenetic tree of the *Fiersviridae* based on the RNA-dependent RNA polymerase gene (alignment length 453 amino acids). The colours correspond to host phyla, green = Streptophyta, blue = Chordata. Viruses from this study are labelled with their phylum and host common name (see Table S1 for details about each host). Branches are scaled according to the number of amino acid substitutions per site, shown in the scale bar. Black circles on nodes show bootstrap support values of more than 90%. The tree is midpoint rooted for display purposes only.

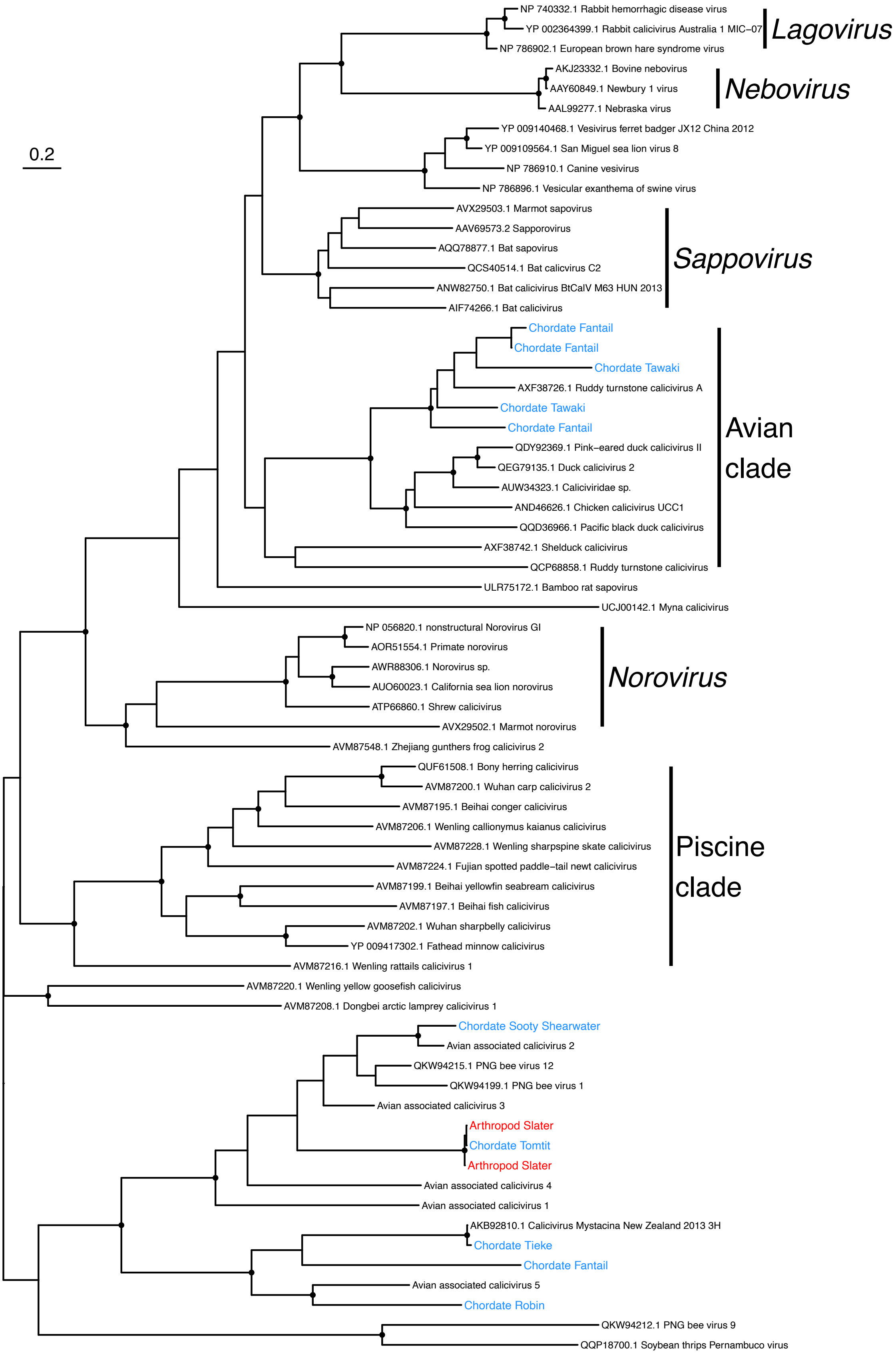
0.2
NP 740332.1 Rabbit hemorrhagic disease virus
YP 002364399.1 Rabbit calicivirus Australia 1 MIC–07
NP 786902.1 European brown hare syndrome virus
Lagovirus
AKJ23332.1 Bovine nebovirus
AAY60849.1 Newbury 1 virus
AAL99277.1 Nebraska virus
Nebovirus
YP 009140468.1 Vesivirus ferret badger JX12 China 2012
YP 009109564.1 San Miguel sea lion virus 8
NP 786910.1 Canine vesivirus
NP 786896.1 Vesicular exanthema of swine virus
AVX29503.1 Marmot sapovirus
AAV69573.2 Sapporovirus
AQQ78877.1 Bat sapovirus
QCS40514.1 Bat calicvirus C2
ANW82750.1 Bat calicivirus BtCalV M63 HUN 2013
AIF74266.1 Bat calicivirus
Sappovirus
Chordate Fantail
Chordate Fantail
Chordate Tawaki
AXF38726.1 Ruddy turnstone calicivirus A
Chordate Tawaki
Chordate Fantail
QDY92369.1 Pink–eared duck calicivirus II
QEG79135.1 Duck calicivirus 2
AUW34323.1 Caliciviridae sp.
AND46626.1 Chicken calicivirus UCC1
QQD36966.1 Pacific black duck calicivirus
AXF38742.1 Shelduck calicivirus
QCP68858.1 Ruddy turnstone calicivirus
Avian clade
ULR75172.1 Bamboo rat sapovirus
UCJ00142.1 Myna calicivirus
NP 056820.1 nonstructural Norovirus GI
AOR51554.1 Primate norovirus
AWR88306.1 Norovirus sp.
AUO60023.1 California sea lion norovirus
ATP66860.1 Shrew calicivirus
AVX29502.1 Marmot norovirus
Norovirus
AVM87548.1 Zhejiang gunthers frog calicivirus 2
QUF61508.1 Bony herring calicivirus
AVM87200.1 Wuhan carp calicivirus 2
AVM87195.1 Beihai conger calicivirus
AVM87206.1 Wenling callionymus kaianus calicivirus
AVM87228.1 Wenling sharpspine skate calicivirus
AVM87224.1 Fujian spotted paddle–tail newt calicivirus
AVM87199.1 Beihai yellowfin seabream calicivirus
AVM87197.1 Beihai fish calicivirus
AVM87202.1 Wuhan sharpbelly calicivirus
YP 009417302.1 Fathead minnow calicivirus
AVM87216.1 Wenling rattails calicivirus 1
Piscine clade
AVM87220.1 Wenling yellow goosefish calicivirus
AVM87208.1 Dongbei arctic lamprey calicivirus 1
Chordate Sooty Shearwater
Avian associated calicivirus 2
QKW94215.1 PNG bee virus 12
QKW94199.1 PNG bee virus 1
Avian associated calicivirus 3
Arthropod Slater
Chordate Tomtit
Arthropod Slater
Avian associated calicivirus 4
Avian associated calicivirus 1
AKB92810.1 Calicivirus Mystacina New Zealand 2013 3H
Chordate Tieke
Chordate Fantail
Avian associated calicivirus 5
Chordate Robin
QKW94212.1 PNG bee virus 9
QQP18700.1 Soybean thrips Pernambuco virus

**Fig. S5**. Phylogenetic tree of the *Caliciviridae* based on the polyprotein gene (alignment length 1313 amino acids). The colours correspond to host phyla, red = Arthropoda, blue = Chordata. Viruses from this study are labelled with their phylum and host common name (see Table S1 for details about each host). Branches are scaled according to the number of amino acid substitutions per site, shown in the scale bar. Black circles on nodes show bootstrap support values of more than 90%. The tree is midpoint rooted for display purposes only.

**Table S1.** Detailed information on each sequencing library. Chordate diet was determined using expert knowledge and published studies.

| Latin name | Library code | Common name | Phylum | Class | Order | Family | Genus | Diet (chordates only) | Samples per library | Total reads | Viral abundance (RPM) | Richness (number of viral families) |
|---|---|---|---|---|---|---|---|---|---|---|---|---|
| *Amphipoda* spp. | AMPH | Landhopper | Arthropoda | Malacostraca | Amphipoda | Talitridae | | | 10 | 110541625 | 48 | 13 |
| *Anthornis melanura* | ANME | Bellbird / korimako | Chordata | Aves | Passeriformes | Meliphagidae | *Anthornis* | Omnivorous | 10 | 229185897 | 9719 | 32 |
| *Apteryx owenii* | APOW | Little spotted kiwi | Chordata | Aves | Apterygiformes | Apterygidae | *Apteryx* | Insectivorous | 3 | 197211236 | 35 | 5 |
| *Araneae* spp. | ARAN | Spider | Arthropoda | Arachnida | Araneae | | | | 10 | 95561385 | 4373 | 29 |
| *Blattodea* spp. | BLAT | Cockroach | Arthropoda | Insecta | Blattodea | Blattidae | *Celatoblatta* | | 5 | 133850527 | 20 | 15 |
| *Poodytes punctatus* | BOPU | Fernbird / mātātā | Chordata | Aves | Passeriformes | Locustellidae | *Poodytes* | Insectivorous | 4 | 189786324 | 14896 | 23 |
| *Coleoptera* spp. | COLE | Beetle | Arthropoda | Insecta | Coleoptera | | | | 10 | 128611425 | 5673 | 25 |
| *Cyanoramphus auriceps* | CYAU | Kakariki / yellow-crowned parakeet | Chordata | Aves | Psittaciformes | Psittaculidae | *Cyanoramphus* | Omnivorous | 6 | 239791291 | 263 | 8 |
| *Dicranoloma billardierei* | DICBIL | Moss | Streptophyta | Bryopsida | Dicranales | Dicranaceae | *Dicranoloma* | | 9 | 150459100 | 16689 | 45 |
| *Diplura* spp. | DIPL | Diplura | Arthropoda | Entognatha | Diplura | | | | 1 | 134084076 | 21 | 7 |
| *Diptera* spp. | DIPT | Fly | Arthropoda | Insecta | Diptera | | | | 10 | 86664024 | 1970 | 14 |
| *Dracophyllum longifolium* | DRALON | Dracophyllum | Streptophyta | Magnoliopsida | Ericales | Ericaceae | *Dracophyllum* | | 10 | 167202609 | 3838 | 35 |

| | | | | | | | | | | | | |
|---|---|---|---|---|---|---|---|---|---|---|---|---|
| *Eudyptes pachyrhynchus* | EUPA | Tawaki / Fiordland crested penguin | Chordata | Aves | Sphenisciformes | Spheniscidae | *Eudyptes* | Piscivorous | 5 | 226372033 | 5458 | 26 |
| *Falco novaeseelandiae* | FANO | New Zealand Falcon / kārearea | Chordata | Aves | Falconiformes | Falconidae | *Falco* | Carnivorous | 1 | 201662827 | 4 | 5 |
| *Gerygone igata* | GEIG | Grey warbler / riroriro | Chordata | Aves | Passeriformes | Acanthizidae | *Gerygone* | Insectivorous | 9 | 226644582 | 407717 | 41 |
| *Geophilomorpha* spp. | GEOP | Centipede | Arthropoda | Chilopoda | Geophilomorpha | | | | 1 | 127720386 | 137 | 13 |
| *Haplotaxida* spp. | HAPL | Pot worm | Annelida | Clitellata | Haplotaxida | Enchytraeidae | | | 4 | 149393198 | 3751 | 15 |
| *Hemiptera* spp. | HEMI | Bug | Arthropoda | Insecta | Hemiptera | | | | 2 | 107380296 | 847 | 11 |
| *Hymenoptera* spp. | HYME | Ant | Arthropoda | Insecta | Hymenoptera | | | | 5 | 99989785 | 23038 | 9 |
| *Isopoda* spp. | ISOP | Slater | Arthropoda | Malacostraca | Isopoda | | | | 10 | 100171201 | 1721 | 16 |
| *Lepidoptera* spp. | LEPI | Moth | Arthropoda | Insecta | Lepidoptera | | | | 2 | 115833281 | 24 | 11 |
| *Lepidothamnus intermedius* | LEPINT | Yellow silver pine | Streptophyta | Pinopsida | Araucariales | Podocarpaceae | *Lepidothamnus* | | 10 | 145809339 | 1302 | 30 |
| *Lithobiomorpha* spp. | LITH | Centipede | Arthropoda | Chilopoda | Lithobiomorpha | | | | 2 | 119449390 | 354 | 20 |
| *Mesostigmata* spp. | MESO | Mite (pred) | Arthropoda | Arachnida | Mesostigmata | | | | 3 | 81062495 | 2089 | 14 |
| *Mohoua novaeseelandiae* | MONO | Brown creeper / pīpipi | Chordata | Aves | Passeriformes | Mohouidae | *Mohoua* | Insectivorous | 4 | 209799281 | 51828 | 28 |
| *Mohoua ochrocephala* | MOOC | Mohua | Chordata | Aves | Passeriformes | Mohouidae | *Mohoua* | Insectivorous | 10 | 187836029 | 12425 | 27 |
| *Nestor meridionalis* | NEME | Kākā | Chordata | Aves | Psittaciformes | Strigopidae | *Nestor* | Omnivorous | 1 | 215076937 | 64 | 7 |
| *Ninox novaeseelandiae* | NINO | Morepork / ruru | Chordata | Aves | Strigiformes | Strigidae | *Ninox* | Insectivorous | 1 | 646268 | 665 | 3 |
| *Nothofagus solandri* | NOTSOL | Mountain beech | Streptophyta | Magnoliopsida | Fagales | Nothofagaceae | *Fuscospora* | | 10 | 124154845 | 2526 | 25 |
| *Oligosoma tekakahu* | OLTE | Te Kakahu/Chalky Island skink | Chordata | Lepidosauria | Squamata | Scincidae | *Oligosoma* | Omnivorous | 10 | 198227057 | 5635 | 20 |
| *Opiliones* spp. | OPIL | Harvestman | Arthropoda | Arachnida | Opiliones | | | | 6 | 102508349 | 1386 | 21 |
| *Opisthopora* spp. | OPIS | Earth worm | Annelida | Clitellata | Opisthopora | | | | 10 | 119919246 | 338 | 10 |

| | | | | | | | | | | | | |
|---|---|---|---|---|---|---|---|---|---|---|---|---|
| *Oribatida* spp. | ORIB | Mite | Arthropoda | Arachnida | Oribatida | | | | 6 | 75305102 | 3091 | 25 |
| *Orthoptera* spp. | ORTH | Weta | Arthropoda | Insecta | Orthoptera | Anostostomatidae | *Hemiandrus* | | 1 | 134022234 | 595 | 5 |
| *Petroica australis* | PEAU | South Island Robin / kakaruai | Chordata | Aves | Passeriformes | Petroicidae | *Petroica* | Insectivorous | 10 | 221633876 | 60066 | 29 |
| *Petroica macrocephala* | PEMA | Tomtit / miromiro | Chordata | Aves | Passeriformes | Petroicidae | *Petroica* | Insectivorous | 5 | 213573349 | 129301 | 34 |
| *Philesturnus carunculatus* | PHCA | Tieke / South Island Saddleback | Chordata | Aves | Passeriformes | Callaeidae | *Philesturnus* | Insectivorous | 10 | 204580700 | 274434 | 30 |
| *Polydesmida* spp. | POLYD | Millipede | Arthropoda | Diplopoda | Polydesmida | | | | 1 | 146389854 | 77 | 12 |
| *Pseudopanax crassifolius* | PSECRA | Lancewood | Streptophyta | Magnoliopsida | Apiales | Araliaceae | *Pseudopanax* | | 9 | 159414298 | 440 | 28 |
| *Pseudoscorpiones* spp. | PSEU | Pseudoscorpion | Arthropoda | Arachnida | Pseudoscorpiones | | | | 6 | 111900992 | 8980 | 36 |
| *Pterodroma inexpectata* | PTIN | Kōrure / Mottled Petrel | Chordata | Aves | Procellariiformes | Procellariidae | *Pterodroma* | Piscivorous | 10 | 242626127 | 14 | 3 |
| *Ardenna grisea* | PUGR | Tītī / Sooty Shearwater | Chordata | Aves | Procellariiformes | Procellariidae | *Ardenna* | Piscivorous | 10 | 195513371 | 195 | 25 |
| *Rhipidura fuliginosa* | RHFU | Fantail / pīwakawaka | Chordata | Aves | Passeriformes | Rhipiduridae | *Rhipidura* | Insectivorous | 10 | 249844003 | 426626 | 31 |
| *Ripogonum scandens* | RIPSCA | Supplejack | Streptophyta | Magnoliopsida | Liliales | Ripogonaceae | *Ripogonum* | | 4 | 139857060 | 500 | 20 |
| *Scolopendromorpha* spp. | SCOL | Centipede | Arthropoda | Chilopoda | Scolopendromorpha | | | | 1 | 86869494 | 41 | 4 |
| *Spirostreptida* spp. | SPIR | Millipede | Arthropoda | Diplopoda | Spirostreptida | | | | 1 | 117989930 | 4911 | 17 |
| *Strigops habroptila* | STHA | Kākāpō | Chordata | Aves | Psittaciformes | Strigopidae | *Strigops* | Herbivorous | 10 | 241609810 | 651 | 8 |
| *Tricladida* spp. | TRIC | Flatworm | Platyhelminthes | Rhabditophora | Tricladida | Geoplanidae | *Australopacifica* | | 1 | 74626559 | 112 | 6 |
| *Vespula germanica* | WASP | German wasp | Arthropoda | Insecta | Hymenoptera | Vespidae | *Vespula* | | 4 | 91075333 | 10299 | 10 |

**Table S2.** Results of permutational analysis of variance (PERMANOVA) models. These models were performed on a Bray-Curtis dissimilarity matrix, created from the OTU abundance table (the abundance of each virus family in each library). Abundance was the number of reads divided by the total reads per library, multiplied by one million (reads per million). Where there were multiple comparisons per model, a pairwise PERMANOVA was used. Significant comparisons (Bonferroni adjusted p-value <0.05) are shown with grey shading.

| Independent variable(s) | Comparison | Degrees of freedom | Sums of squares | F model | $R^2$ | p-value | adjusted p-value |
|---|---|---|---|---|---|---|---|
| Host phyla | Main effect | 4 | 2.99 | 1.68 | 0.13 | 0.0001 | 0.0001 |
| Host phyla | Chordata vs Arthropoda | 1 | 0.82 | 1.79 | 0.04 | 0.003 | 0.03 |
| | Chordata vs Streptophyta | 1 | 1.16 | 2.73 | 0.11 | 0.001 | 0.01 |
| | Chordata vs Annelida | 1 | 0.66 | 1.50 | 0.07 | 0.03 | 0.3 |
| | Chordata vs Platyhelminthes | 1 | 0.54 | 1.23 | 0.06 | 0.05 | 0.5 |
| | Arthropoda vs Streptophyta | 1 | 0.92 | 2.03 | 0.08 | 0.001 | 0.01 |
| | Arthropoda vs Annelida | 1 | 0.56 | 1.19 | 0.05 | 0.05 | 0.5 |
| | Arthropoda vs Platyhelminthes | 1 | 0.48 | 1.00 | 0.05 | 0.5 | 1 |
| | Streptophyta vs Annelida | 1 | 0.77 | 2.15 | 0.26 | 0.04 | 0.4 |
| | Streptophyta vs Platyhelminthes | 1 | 0.59 | 1.63 | 0.25 | 0.1 | 1 |
| | Annelida vs Platyhelminthes | 1 | 0.55 | 1.53 | 0.61 | 0.3 | 1 |
| Host class | Main effect | 12 | 6.75 | 1.28 | 0.30 | 0.0001 | 0.0001 |
| Host order | Main effect | 36 | 18.21 | 1.37 | 0.80 | 0.0001 | 0.0001 |

| | | | | | | | |
|---|---|---|---|---|---|---|---|
| Host order + host diet (chordates only) | Insectivore vs non-insectivore | 1 | 0.49 | 1.54 | 0.06 | 0.01 | 0.03 |
| | Plant-eater vs non-plant-eater | 1 | 0.56 | 1.75 | 0.07 | 0.0008 | 0.002 |
| Module | 4 vs 3 | 1 | 0.78 | 2.87 | 0.12 | 0.002 | 0.012 |
| | 4 vs 1 | 1 | 1.66 | 6.21 | 0.16 | 0.001 | 0.006 |
| | 4 vs 2 | 1 | 1.10 | 4.74 | 0.18 | 0.001 | 0.006 |
| | 3 vs 1 | 1 | 0.98 | 3.56 | 0.13 | 0.001 | 0.006 |
| | 3 vs 2 | 1 | 0.90 | 4.13 | 0.24 | 0.001 | 0.006 |
| | 1 vs 2 | 1 | 1.37 | 5.79 | 0.19 | 0.001 | 0.006 |

**Table S3**. The mean (± standard deviation) of permutational analysis of variance (PERMANOVA) models over 100 rarefied data sets of the original OTU abundance table, with module number as the independent variable. Abundance was the natural log of the raw abundances.

| **Comparison** | **Degrees of freedom** | **Mean Sums of squares (±SD)** | **Mean F model (±SD)** | **Mean R2 (±SD)** | **mean p-value (±SD)** | **mean adjusted p-value (±SD)** |
|---|---|---|---|---|---|---|
| 4 vs 3 | 1 | 0.80 (0.04) | 2.65 (0.15) | 0.11 (0.005) | 0.006 (0.003) | 0.04 (0.02) |
| 4 vs 1 | 1 | 1.61 (0.06) | 4.90 (0.19) | 0.13 (0.004) | 0.001 (0) | 0.006 (0) |
| 4 vs 2 | 1 | 1.84 (0.06) | 6.83 (0.28) | 0.24 (0.007) | 0.001 (0.0001) | 0.006 (0.0006) |
| 3 vs 1 | 1 | 0.90 (0.06) | 2.47 (0.13) | 0.01 (0.005) | 0.002 (0.001) | 0.01 (0.007) |
| 3 vs 2 | 1 | 1.02 (0.04) | 3.47 (0.21) | 0.21 (0.01) | 0.002 (0.0008) | 0.01 (0.005) |
| 1 vs 2 | 1 | 1.41 (0.04) | 4.20 (0.18) | 0.15 (0.005) | 0.001 (0.0001) | 0.006 (0.0008) |

.

**Table S4**. The hosts and viruses belonging to each module, as identified by the modularity analysis. Shading represents the four different modules and corresponds to the same colours used in Figure 3.

| **Module** | **Hosts (library code)** | **Virus families** |
|---|---|---|
| Module 1 | COLE | *Adenoviridae* |
| | HEMI | *Adintoviridae* |
| | GEOP | *Aliusviridae* |
| | LITH | *Artoviridae* |
| | BLAT | *Baculoviridae* |
| | DIPL | *Benyviridae* |
| | OPIS | *Chuviridae* |
| | TRIC | *Circoviridae* |
| | DIPT | *Euroniviridae* |
| | OPIL | *Iridoviridae* |
| | AMPH | *Lispiviridae* |
| | POLYD | *Metaviridae* |
| | SPIR | *Mimiviridae* |
| | ORIB | *Mononiviridae* |
| | LEPI | *Nairoviridae* |
| | HAPL | *Nudiviridae* |
| | ARAN | *Nyamiviridae* |
| | ORTH | *Orthomyxoviridae* |

| | | |
|---|---|---|
| | | *Parvoviridae* |
| | | *Peribunyaviridae* |
| | | *Phasmaviridae* |
| | | *Phenuiviridae* |
| | | *Polyomaviridae* |
| | | *Poxviridae* |
| | | *Rhabdoviridae* |
| | | *Tospoviridae* |
| | | *Totiviridae* |
| | | *Unclassified Martellivirales* |
| | | *Unclassified Mononegavirales* |
| | | *Xinmoviridae* |
| Module 2 | DRALON | *Amalgaviridae* |
| | PSECRA | *Arenaviridae* |
| | MESO | *Aspiviridae* |
| | DICBIL | *Barnaviridae* |
| | NOTSOL | *Betaflexiviridae* |
| | PSEU | *Birnaviridae* |
| | RIPSCA | *Botourmiaviridae* |
| | LEPINT | *Bromoviridae* |
| | | *Caulimoviridae* |
| | | *Chrysoviridae* |

| | |
|---|---|
| | *Closteroviridae* |
| | *Deltaflexiviridae* |
| | *Endornaviridae* |
| | *Fusariviridae* |
| | *Geminiviridae* |
| | *Genomoviridae* |
| | *Herpesviridae* |
| | *Hypoviridae* |
| | *Kitaviridae* |
| | *Megabirnaviridae* |
| | *Mitoviridae* |
| | *Mymonaviridae* |
| | *Narnaviridae* |
| | *Partitiviridae* |
| | *Phycodnaviridae* |
| | *Pithoviridae* |
| | *Polymycoviridae* |
| | *Potyviridae* |
| | *Qinviridae* |
| | *Rountreeviridae* |
| | *Tectiviridae* |
| | *Togaviridae* |

| | | |
|---|---|---|
| | | *Unclassified Bunyavirales* |
| | | *Unclassified Tymovirales* |
| | | *Virgaviridae* |
| | | *Yueviridae* |
| Module 3 | FANO | *Atkinsviridae* |
| | NEME | *Fiersviridae* |
| | STHA | *Hepadnaviridae* |
| | OLTE | *Herelleviridae* |
| | ISOP | *Inoviridae* |
| | PUGR | *Leviviridae* |
| | EUPA | *Marnaviridae* |
| | | *Microviridae* |
| | | *Myoviridae* |
| | | *Nimaviridae* |
| | | *Picobirnaviridae* |
| | | *Podoviridae* |
| | | *Sarthroviridae* |
| | | *Siphoviridae* |
| | | *Steitzviridae* |
| | | *Unclassified Caudovirales* |
| | | *Unclassified Ortervirales* |
| Module 4 | HYME | *Alphatetraviridae* |

| | |
|---|---|
| ANME | *Anelloviridae* |
| MONO | *Astroviridae* |
| SCOL | *Autographiviridae* |
| RHFU | *Caliciviridae* |
| BOPU | *Carmotetraviridae* |
| WASP | *Dicistroviridae* |
| GEIG | *Flaviviridae* |
| CYAU | *Hepeviridae* |
| APOW | *Iflaviridae* |
| MOOC | *Luteoviridae* |
| NINO | *Mayoviridae* |
| PTIN | *Mesoniviridae* |
| PEAU | *Nodaviridae* |
| PHCA | *Permutotetraviridae* |
| PEMA | *Picornaviridae* |
| | *Polycipiviridae* |
| | *Polydnaviridae* |
| | *Reoviridae* |
| | *Retroviridae* |
| | *Schitoviridae* |
| | *Secoviridae* |
| | *Sinhaliviridae* |

| | |
|---|---|
| | *Solemoviridae* |
| | *Solinviviridae* |
| | *Tombusviridae* |
| | *Tymoviridae* |
| | *Unclassified Nidovirales* |
| | *Unclassified Picornavirales* |

**Table S5**. Details of the sequence alignments used to estimate the phylogenetic trees for each of the four viral families chosen for phylogenetic analysis out of a total of 112 viral families. The % pairwise identity refers to the percentage of pairwise residues that are identical in the alignment, excluding gap-gap residues.

| **Family** | **Protein** | **Alignment length (amino acids)** | **Number of sequences** | **% Pairwise Identity** |
|---|---|---|---|---|
| *Parvoviridae* | Non-structural protein 1 | 584 | 292 | 15.8 |
| *Caulimoviridae* | Polyprotein | 938 | 123 | 40.9 |
| *Fiersviridae* | RNA-dependent RNA polymerase | 453 | 336 | 32.2 |
| *Caliciviridae* | Polyprotein | 1111 | 68 | 20.1 |

**Table S6**. The abundance (expressed as reads per million) and percentage abundance of viruses from each host phylum for one viral family per module.

| **Host phylum** | ***Parvoviridae* (module 1)** | ***Caulimoviridae* (module 2)** | ***Fiersviridae* (module 3)** | ***Caliciviridae* (module 4)** |
|---|---|---|---|---|
| Arthropoda | 351 (99%) | 0 (0%) | 0 (0%) | 1 (0.03%) |
| Chordata | 2 (0.6%) | 2 (0.2%) | 10 (99%) | 3952 (99.98%) |
| Streptophyta | 0.1 (0.03%) | 918 (99.8%) | 0.1 (1%) | 0 (0%) |
| Annelida | 3 (0.3%) | 0 (0%) | 0 (0%) | 0 (0%) |
| Platyhelminthes | 0 (0%) | 0 | 0 (0%) | 0 (0%) |

**Data S1. (Separate file, OTU_table_raw_abundances.xlsx)**

The operational taxonomic unit table of raw abundances used in the beta diversity analysis and unipartite network generation and analysis, with abundance corrected to reads per million (raw abundance divided by the total number of reads, multiplied by one million). This table was also used for the module analysis, with abundance corrected to log abundance.
